# eDNA provides accurate population abundance estimates with bioenergetics and particle mass-balance modelling

**DOI:** 10.1101/2024.12.23.629923

**Authors:** Julien Beaulieu, Matthew C. Yates, Dylan J. Fraser, Melania E. Cristescu, Alison M. Derry

## Abstract

Anthropogenic activities have led to an unprecedented crisis in freshwater biodiversity loss. The capacity to monitor the abundance of wild populations is critical to conserving biodiversity, but conventional physical specimen collection methods are invasive, costly, and labour-intensive. Environmental DNA (eDNA) offers a promising alternative, being easy to sample, with studies under controlled laboratory conditions showing consistent correlations between eDNA concentration and abundance. However, applying eDNA to monitor abundance remains contentious, as eDNA particle dynamics and the ecology of eDNA production can decouple this relationship in natural ecosystems. To address this, we provide a novel modeling method to produce population estimates from eDNA. We integrated bioenergetics and mass-balance frameworks to relate eDNA concentrations to freshwater fish population abundance estimated through conventional mark-recapture in Brook Trout (Salvelinus fontinalis) across nine Rocky Mountains lakes, five of which underwent size-selective harvesting over two years. Our integrated framework improved the variance explained in eDNA concentrations from 24% to 71%. The integrated model accurately distinguished most (94%) abundance estimates across populations and sampling periods, detecting both natural and harvest-induced reductions in abundance within several populations. This study is the first to empirically integrate the DNA production mechanism and particle dynamics and provide a new methodological approach enabling rapid and accurate abundance quantification. We also discuss how this new tool can be integrated in existing monitoring programs.

## Introduction

Anthropogenic activities have triggered an unparalleled biodiversity crisis in freshwater ecosystems (Albert et al., 2021). Conserving remaining biodiversity, remediating degraded ecosystems, and halting the decline of endangered populations demand rigorous monitoring and proactive aquatic ecosystem management (Ho & Goethals, 2019). However, freshwater ecosystems are notoriously difficult to monitor, with the decline in freshwater fish populations often referred to as an ‘invisible collapse’ (Howarth et al., 2023; Post et al., 2002). Monitoring the abundance and biomass of species in aquatic ecosystems can be particularly challenging, due to its labour- and resource-intensive nature. These challenges are magnified at larger geographic scales, where obtaining such data for remote or large freshwater waterbodies can be exceedingly difficult if not impossible. As a result, the availability of such critical data for conservation and management is often severely limited.

The analysis of environmental DNA (eDNA), which refers to DNA particles found in environmental mediums such as air, soil, and water (Ficetola et al., 2008), has revolutionised biomonitoring in natural ecosystems. The analysis of eDNA has proven highly effective for quantifying ecosystem-level biodiversity as well as detecting cryptic, rare, and traditionally difficult-to-detect species in freshwater ecosystems (Jerde, 2019; Sepulveda et al., 2020). However, numerous studies have demonstrated consistent and significant positive correlations with the concentration of eDNA and the abundance or biomass of organisms within aquatic environments; these studies have demonstrated its strong potential for providing estimates of population abundance and/or biomass in natural ecosystems (Rourke et al., 2022; Yates et al., 2019). Yet, the biological processes involved in the production of eDNA production are complex and, as a physical particle, numerous environmental parameters can impact its degradation and transportation in natural ecosystems (Carraro et al., 2018; Stewart, 2019). Collectively, these processes can decouple the relationships between the concentration of eDNA in natural ecosystems and organism abundance/biomass (Yates, Cristescu, et al., 2021). This can be readily observed in the discrepancy between the strong correlation eDNA exhibits with abundance under controlled laboratory conditions relative to the moderate correlations observed in studies conducted within natural ecosystems (Yates et al., 2019).

Our understanding of the behaviour and dynamics of eDNA particles in natural ecosystems is constantly growing and improving through rigorous research occurring in labs across the globe. The decay rate of eDNA, for example, can depend on complex interactions among abiotic conditions (e.g., temperature, pH and UV), biotic conditions (e.g., bacterial activity), and eDNA properties (amplicon and particle size) (Brandão-Dias et al., 2023; T. Jo et al., 2019; T. Jo & Minamoto, 2021; Lamb et al., 2022; Saito & Doi, 2021a; Strickler et al., 2015). In both lentic and lotic aquatic ecosystems, eDNA particles move within water masses at different rates, which can additionally affect their deposition. The concentration measured at one place can thus be influenced by eDNA production upstream and/or its movement within currents or within thermocline layers (Jane et al., 2015; Littlefair, Rennie, et al., 2021; Pont, 2024; Shogren et al., 2017).

Metabolic and bioenergetic theory has also recently been integrated as a framework to relate eDNA production to population density and biomass by accounting for the physiological processes involved in its production (Yates et al., 2023). Metabolic processes, for example, scale allometrically with organism size (Allegier et al., 2015; Vanni & McIntyre, 2016). The strong link between eDNA production and organism metabolism implies that eDNA production likely scales allometrically within and across species (Yates et al., 2023; Yates, Wilcox, et al., 2021). Similarly, temperature has long been recognized as having a strong impact upon physiological rates within poikilothermic fishes (Thornton & Lessem, 1978), which similarly extends to eDNA production (Caza-Allard et al., 2022; T. Jo et al., 2019). As a result, eDNA production can be modelled as a function of allometrically scaled organismal mass weighted with a temperature-dependent function to adjust for the impact of the metabolic rate of the organism on eDNA production (Deslauriers et al., 2017; Hanson et al., 1997; Yates, Cristescu, et al., 2021; Yates, Wilcox, et al., 2021), e.g.:

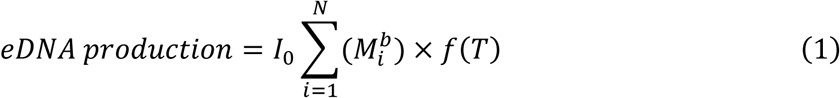

Where *I_0_* is the eDNA production coefficient, *N* is the number of individuals in the population, *M_i_* is the mass of the i^th^ individual, *b* is the allometric scaling coefficient, and *f(T)* is the temperature-dependent function, which considers the maximal metabolic rate of the organism at a specific temperature (Yates, Cristescu, et al., 2021).

The pseudo-steady-state concentration of eDNA in natural ecosystems is a product of complex dynamics between eDNA production, degradation, and transportation. Integrating across these processes provides an understanding of dynamics that affect particle input and output rates; as a result, the application of a mass-balance approach is ideally suited to modelling eDNA steady-state concentrations, in a similar manner as classic models used for estimating phosphorus concentration (Chapra & Reckhow, 1983; T. Jo et al., 2019; Sassoubre et al., 2016). If we assume the concentration of eDNA to be constant in time, we can state:

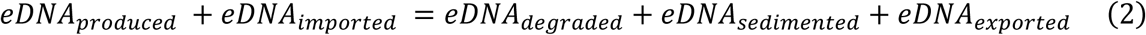

Where the eDNA production rate is given by the metabolic/bioenergetics function described above (equation 1), and eDNA imported describes the rate eDNA enters the system from up-stream sources. The rate at which eDNA is degraded is typically described by an exponential decay curve as a function of temperature, pH, and other factors that can influence eDNA degradation rate, with ‘*k’* being the degradation constant (Kagzi et al., 2022). No research has yet examined eDNA sedimentation rates independent of particle decay, but sedimentation of particles in the water are also typically modeled using exponential decay curves (Chapra & Reckhow, 1983). The rate at which eDNA is exported (i.e., transported) refers to the amount of eDNA exiting by the outflow of a water body, which can be calculated as the outflow times the concentration at the surface (Chapra & Reckhow, 1983). By developing equation 2 and isolating eDNA concentration, the formula can be expressed as:

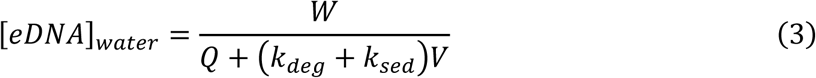

Where *[eDNA]_water_* is the eDNA concentration in the water, *W* is the average eDNA production (equation 1) plus the average eDNA importation from upstream, *V* is the volume of the water body, *Q* is the water flow rate, *K_deg_* is the degradation constant at these specific environmental conditions, and *K_sed_* is the sedimentation rate.

Modelling that integrates both bioenergetics and mass balance approaches to eDNA concentrations to estimate total population abundance and biomass in natural ecosystems remains unexplored. Yet, it holds considerable potential for improving the capacity to link eDNA concentrations observed in natural ecosystems to wild population abundance. With refined quantitative knowledge on eDNA production, decay, and transport rates, eDNA could become a cost-effective tool for species quantification in aquatic systems (Huang et al., 2022; Yates, Cristescu, et al., 2021). To date, however, no study has integrated across all these processes to generate total population abundance/biomass estimates in natural ecosystems.

We investigated the application of an integrated bioenergetics and mass-balance approach to estimate population abundance from eDNA concentrations observed in natural lentic aquatic ecosystems. Our aims were to: i) differentiate total abundance *across* populations; and ii) detect changes in total abundance *within* populations. To accomplish these objectives, we quantified Brook Trout (*Salvelinus fontinalis*) eDNA concentrations and total abundance (using mark-recapture techniques) in nine Rocky Mountain lakes (CA) across two seasons for two years. This was done while simultaneously conducting an annual experimental size-selective harvest in five of the nine lakes to manipulate total abundance and population density across the study period. We then investigated the added power of an integrated bioenergetics and mass-balance approach in explaining observed eDNA concentrations compared to approaches previously taken in the literature. This included: i) correlating eDNA concentrations as a linear function of population abundance, and ii) correlating eDNA concentrations as a function of allometrically scaled population biomass. We predicted that taking an integrated bioenergetic and mass-balance approach that holistically accounts for the processes involved in eDNA production, degradation, and transportation would increase the power of eDNA to detect changes in the abundance and biomass of Brook Trout across space and time.

## Methods

### Study system

Our study included nine headwater mountain lakes in the Canadian Rockies, located within Banff, Yoho, and Kootenay National Parks (Canada; Fig. 1). The nine study lakes contain fish communities largely dominated by Brook Trout (*Salvelinus fontinalis*) that were introduced to the lakes between the 1920s and the 1970s and are now invasive in the region (Messner et al., 2013). Six of the lakes were solely inhabited by Brook Trout, while the three other had fish communities also containing one to five other species. Five of the nine study lakes were subjected to a size-selective harvest treatment where approximately 64 % of each population was removed each year during Fall from 2017 to 2019, targeting the largest-bodied individuals for harvest. At the beginning of the experiment (2017), the Brook Trout populations differed in density and population size structure between lakes, and the harvesting treatment was successful at inducing changes in population demography over the duration of the three-year experimental manipulation (Matte et al., 2023). The other four study lakes were monitored as references in which Brook Trout populations were not directly manipulated (Appendix 1). All lakes are small, cold, oligotrophic headwater lakes, but they span a 1000 m elevation gradient over which there are differences between lakes in temperature, pH, and nutrient concentrations (Appendix 1; Trépanier-Leroux et al., 2023).

**Figure 1.**
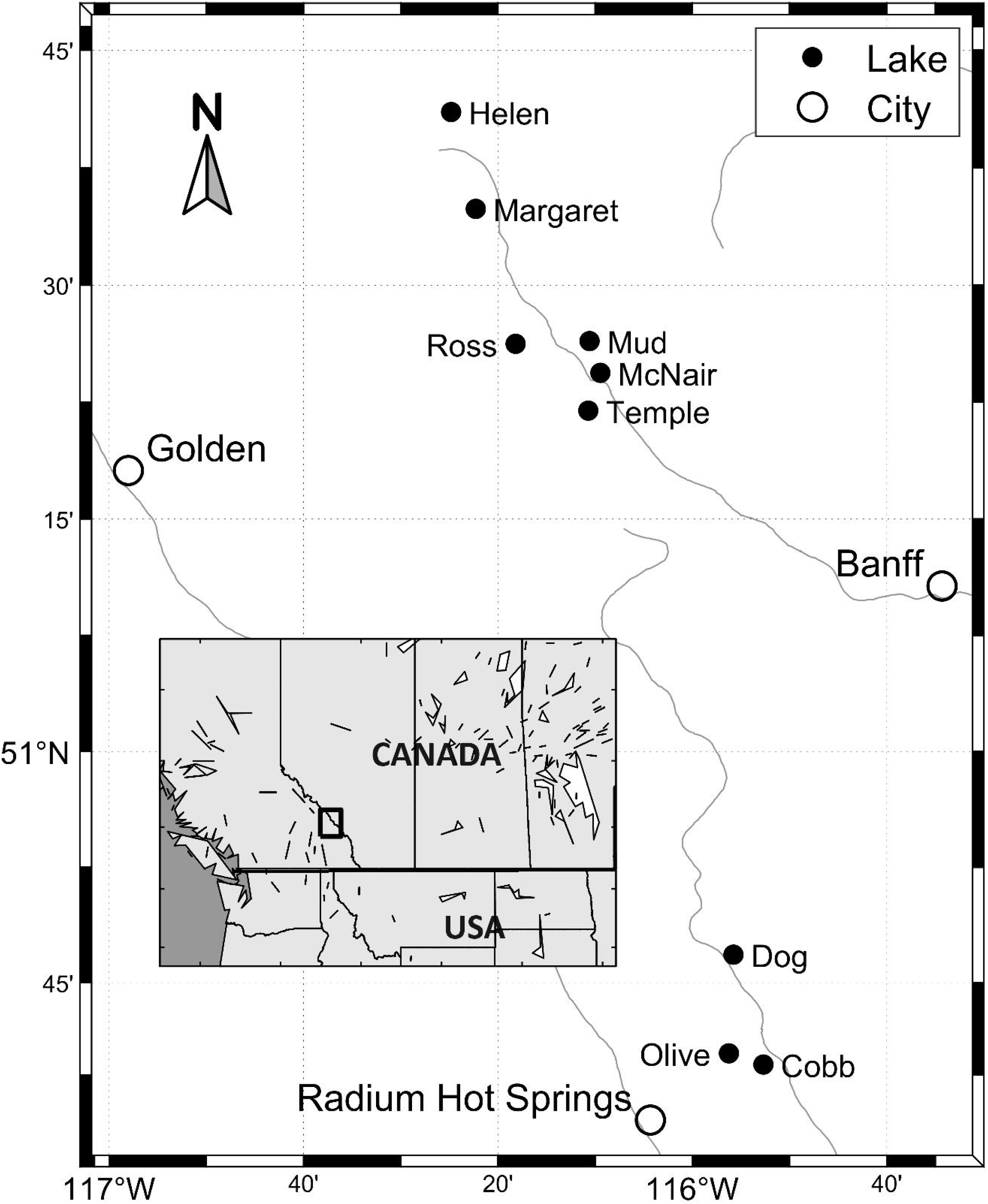
Map of the study area showing the locations of the lakes included in the research.

### Field sampling of eDNA

Environmental DNA samples were collected each year between 2018 and 2019 for two time periods: i) in Summer before harvesting (June/July); and ii) in the Fall (September) following harvest activities. A systematic stratified survey design was used to collect eDNA samples, in which 4 samples each were collected from the pelagic and littoral zones of each study lake. In the littoral zones, eDNA sample sites were distributed equidistantly around the perimeter of the lake. Littoral samples were collected approximatively 1-3 m from the shore at a minimum depth of 30 cm from the surface, but approximately 15 cm above the lake bottom. In the pelagic zones, four samples were collected at equidistant intervals across a transect that followed the longest axis of the lake and went through the lake center (defined as the midpoint of the longest axis). Pelagic samples were collected at a depth of 0.5 to 1 m from the surface. Water samples were collected with 700 ml Whirl-Pak™ bags from a canoe assigned to sample a single specific lake, or from an inflatable kayak, which was decontaminated 48 hours prior with a 2% regular strength household bleach solution for 15 min (including paddle and lifejacket) and left to dry in the sun.

Water samples were immediately filtered on the lakeshore on a water filtering manifold (Wildco, Florida, USA) that was decontaminated with a 20-30% bleach solution 2-12 h prior to sampling. Manifold components were stored individually in plastic zippered bags after bleaching and during transportation. All samples were stored in the shade prior to sampling, and all filtering was conducted in the shade under a tarp. One litre of water from each of the eight water sampling sites per lake was filtered through a 0.7 μm glass fibre filter (GE Healthcare Life Sciences, Ontario, Canada) using a vacuum hand pump (Soil Moisture, California, USA). Vacuum pumps were decontaminated between lakes by wiping with a 30% bleach solution and resting for 10 minutes, after which they were wiped with distilled water to remove residual bleach. All littoral samples were filtered on one manifold, and all pelagic samples were filtered on another to prevent cross-contamination between samples from different lake zones. As a negative control, 500 mL of distilled water was filtered prior to filtering lake samples. After filtration, filters were folded using metal forceps (decontaminated using a 30% bleach solution for 15 minutes) and stored in a sterile 2 mL microcentrifuge tube containing 700 μL of ATL buffer that was then individually placed in a plastic zippered bag and stored in a cooler bag that was decontaminated by wiping with a 30% bleach solution. If a filter became clogged (i.e. < 1 L of water was filtered), the final volume of water filtered was recorded and the sample was then stored in buffer. Samples were then transported in a cooler bag with bleached frozen gel packs to Kootenay Crossing (BC, Canada), where they were placed in a second zippered plastic bag and stored in a -20 ℃ freezer. At the end of the sampling season, all samples were transported to Montreal on dry ice and placed in a -80 ℃ freezer until extraction. All protocols for contamination prevention in the field, as well as during transport and storage can be found in Appendix 2.

### Laboratory analysis of eDNA

DNA was extracted from each filter using Qiagen DNeasy Blood and Tissue^TM^ kit and Qiashredder^TM^ columns following a modified extraction protocol (see Yates et al. (2021a) for more details). Extracted DNA was eluted into 130 μL of AE buffer and stored in a clean -20 ℃ freezer solely dedicated to the storage of extracted eDNA product (i.e. no post-PCR products or tissue samples). An extraction blank of 700 μL of ATL buffer was included in all batches as a control. Methods for contamination prevention are detailed in Supplementary Information (Appendix 2).

Brook Trout eDNA concentration was quantified using quantitative PCR (qPCR) with the *cyt b* TaqMan minor groove assay (BRK2) described in (Wilcox et al., 2013). For details on PCR reaction preparation, component concentrations, standard curve preparation, and cycling conditions, see (Yates, Glaser, et al., 2021). All samples were run in triplicate in a 20 μL reaction volume with 5 μL of template DNA on a Stratagene MX 3000P thermal cycler. Reactions were spiked with an internal positive control (IPC) to test for inhibition, which was defined as a 1 > Ct shift in the IPC relative to a reaction containing 5 μL of nuclease-free ultrapure water. If inhibition was detected, samples were diluted down to 3 μL and re-run. Brook Trout eDNA concentrations at each site were calculated by averaging replicate values, and final copy number data were converted to eDNA concentration per L of water filtered (copies/L). Standard curve *R^2^* values for the qPCR plates ranged from 0.98-0.99, and reaction efficiency varied between 83.1 and 98.6%. The efficiency of two of the 11 plates was suboptimal (< 85%). However, data generated from these two plates were retained, as their inclusion did not impact results (Appendix 3). Samples associated with negative controls that contained more than 5 copies/rx (on average) were removed from the analysis due to contamination concerns (Cobb Lake samples from Fall 2019). Most samples collected across lakes per sampling period (26/36 sampling events) showed no contamination, while the remaining contamination in field controls was low enough to not impact the estimated lake eDNA concentrations.

### Brook Trout mark-recapture and population size structure estimation

Mark-recapture studies of Brook Trout were conducted in all lakes during May and June in both 2018 and 2019 (Matte et al., 2023; Yates, Glaser, et al., 2021). The exception was Cobb Lake where, in addition to mark-recapture sampling in May/June, a series of isolated marking events continued until September for both years. Fish were captured using a combination of fyke nets, angling, and backpack electrofishing. Large (1 m hoop diameter, 2 cm mesh) and small (0.7 m hoop diameter and 0.8 cm mesh) fyke nets were distributed around the perimeter of lakes with the lead attached to shore and the end of the trap facing the center of the lake. Nets were checked daily. Angling was used to supplement Brook Trout capture efforts at sites where fyke catchability was low (predominantly Cobb Lake). Marks were also assigned to Brook Trout captured by electrofishing the shore and inlets/outlets of lakes.

Captured Brook Trout were anesthetized using clove oil and measured for fork length (± 1 mm) and mass (± 0.1 g). Any unmarked Brook Trout were gastrically tagged with a BioMark HPT8 pre-loaded Passive Integrated Transponder (PIT) tag (Boise, Idaho, USA). Only Brook Trout greater than or equal to 80 mm were tagged to reduce tagging mortality. Other species, when present, were not tagged. The tag number of any recaptured Brook Trout was recorded. Recovered fish were released in the center of the lake to standardize release location and promote mixing. Marking ceased once recapture ratios approached twenty five percent for several consecutive days to standardize marking efforts across all populations and to ensure that enough Brook Trout were tagged to facilitate estimation of census size (Nc) with credible intervals within 10% to 25% of true values, following general methodologies reviewed in (Krebs, 2009; Matte et al., 2023).

To obtain representative snapshots of the size structure of each Brook Trout population before and following harvest, population size structure estimates were conducted in August 2018 and 2019 (Matte et al., 2023). The exception was Cobb Lake where size structure assessments continued to October 12^th^. During mark-recapture, fyke-nets were deployed in littoral zone areas extending to the centre of the lake and, as a result, size-structure assessments obtained from fyke nets may be more biased towards small-medium bodied individuals that tend to prefer littoral habitats (Tiberti et al., 2017). To obtain a relatively unbiased estimate of population size structure, Brook Trout were captured in large and small sinking mixed mesh gillnets with clear monofilament distributed throughout the lake. Large mixed-mesh gillnets were 15.6 m long, 1.8 m deep and had an equal area of 64-51-89-38-76 mm mesh panels. Small mixed-mesh gillnets were 12.5 meters long, 1.8 meters deep, and consisted of an equal area of 32-19-38-13-25 mm mesh panels. Such index nets are widely used in North America for size structure assessments (Bonar et al., 2009; Hubert et al., 2012; Johnson, 1983; Post et al., 1999; Ward et al., 2012), as these capture a representative size/age structure of fish populations (Morgan, 2002). Nets were checked daily and moved to different locations across the lake, to capture a representative sample of Brook Trout in each lake. Sampling ceased when approximately five to ten percent of the population was captured, apart from Cobb Lake where the harvested fish were also included for size structure assessment such that approximately 71% of individuals were used. Captured Brook Trout were euthanized with clove oil, PIT tags were recorded, and length/mass data were collected as described for the marking period.

Schnabel population abundance estimates were used to estimate the abundance of Brook Trout in a lake (Matte et al., 2023; Schnabel, 1938). All size structure assessment removals were pooled together into one final sampling event for the population abundances which controlled for the removal of marks at large. Note that abundances only account for Brook Trout greater than the minimum tagging size (80 mm fork length), which correspond to all individuals age 1 and older (Matte et al., 2023). All population abundace estimates were conducted in R (CoreTeam, 2020) with the mrClosed function from the Fisheries Stock Assessment package FSA (Ogle, 2015). Credible intervals for Schnabel population abundance estimates followed recommendations from (Seber, 2002) as implemented in the FSA package.

### Size-selective harvesting of Brook Trout

Populations at Cobb, Mud, Olive, Ross, and Temple lakes were harvested annually with size-selective gillnets (25– 89 mm mesh) targeting the top 75% (but effectively harvesting the top 64% on average) of individual body sizes according to the size distributions estimated during stock assessment (i.e. depleting the largest size classes sequentially until the only remaining individuals were in the lowest 25% quartile of the original size structure). Details can be found in Matte et al. (2023).

### Lake temperature and its effect on the physiology of eDNA production

To adjust for maximal metabolic rate of fish at a given temperature, we used a temperature-dependent function specific to Brook Trout published from previous empirical studies (Deslauriers et al., 2017; Hartman & Cox, 2008). Temperature data were constituted of both high frequency (15 min) surface temperature from data loggers (Onset Hobo®; UA-002-64, Onset Computer Corporation, Cape Cod, Massachusetts) and vertical profiles from a YSI multiparameter sonde (model 10,102,030; YSI Inc.). The dataloggers were deployed at 0.5 m from the surface in June and retrieved in September. The YSI vertical profiles were taken at the deepest point of the lake at the same time as eDNA sampling, except six lakes in Fall 2019 where a YSI sonde malfunction necessitated us to use data from the same time period in 2018 as a proxy. There was an approximately +/- 2°C difference between Fall 2018 and 2019 for the lakes that had YSI data for both seasons and as a result, this likely had a negligible impact on model estimates. When possible, mean surface temperature over the interval from two weeks before to two weeks after each eDNA sampling event per lake was used as input in the temperature-dependent function in the eDNA mass balance model. However, due to logistic constraints and material failures, some lakes and sampling periods had temperature data for only a subset of the time intervals or only the epilimnion mean temperature from the YSI vertical profiles. A table with the lakes, dates, and temperature data can be found in the supplementary materials (Appendix 4).

### Lake hydrology

To calculate the rate of lake outflow, lake watershed area was estimated using the flow direction and flow accumulation hydrologic models from ArcGIS pro V.3.2. The input data were 15 m resolution Digital Elevation Models (DEM) for 7 out of the 9 study lakes. However, the 15 m resolution was too low to effectively determine the watershed for two of the lakes (Mud and McNair). Fifteen m resolution DEM are publicly available form the Government of Canada and under Open Government Licence (https://open.canada.ca/en/open-government-licence-canada). The Mud and McNair lakes required high resolution data, but none were available for their area. The Mud Lake watershed area was therefore estimated by process of elimination by estimating the watershed area of the surrounding larger water courses. The remaining area that was not part of any other watershed was assumed to be Mud Lake’s watershed. We could not obtain watershed area for McNair Lake. This missing data was handled with data imputation before fitting using the mice package (Buuren & Groothuis-Oudshoorn, 2011). The lake outflow rate was estimated by multiplying each lake watershed area with the flux per area of watershed from hydrological stations from the same area (Kootenay: Kootenay River: 50.886944, -116.045833; Banff: Pipestone River: 51.433056, -116.174722; Environment and Natural Resources Canada, 2024). The rate of outflow for each lake at each sampling period was calculated as the mean outflow from two weeks before the sampling date to two weeks after.

### Lake volume and epilimnion volume

Detailed bathymetric maps and lake volume produced by high frequency sonar were available for only four study lakes (Temple, Olive, Helen, and Margaret). For the remaining five study lakes, we used older bathymetric maps from Parks Canada that were georeferenced and digitised in ArcGIS Pro. The depth between isobaths were interpolated using Bayesian Kriging method. The epilimnion volume was taken as the input in the mass balance model (equation 5 and 8). Epilimnetic volume was chosen because eDNA samples were collected from the epilimnion of each lake; this was done because most study lakes undergo thermal stratification during Summer, and because Brook Trout spend most of their time feeding and resting in surface waters (Tiberti et al., 2017). Additionally, research has demonstrated that the transfer of eDNA across the thermocline is limited during lake stratification (Littlefair, Hrenchuk, et al., 2021). The epilimnion volume was calculated as the mean thermocline depth times the lake surface area from the bathymetric maps. The mean thermocline depth was calculated from two YSI sonde vertical temperature profiles per Summer. For the study lakes that were not thermally stratified (assessed from the YSI vertical temperature profile) at the time of eDNA sampling period, total lake volume (estimated from the bathymetric maps) was used as input in mass-balance models for that time point because eDNA signal is expected to become homogenous throughout the water column during mixing (Littlefair, Hrenchuk, et al., 2021). The extent of thermal stratification during the Fall 2019 could not be quantified due to the malfunction of the YSI sonde. Lakes that were mixed in Fall 2018 were therefore also assumed to be mixing in Fall 2019 except for Margaret Lake, where stratification was very weak in Fall 2018 and it was not fully mixed. As Fall 2019 was colder than Fall 2018, we assumed that Margaret Lake was mixing in Fall 2019; this increased model fit for this lake.

### Statistical analyses

#### Constructing and evaluating an integrated bioenergetics/mass-balance model

To obtain a representative mean eDNA concentration for each lake and sampling period, the littoral and pelagic samples were weighted according to the littoral and pelagic zone area in each lake. The littoral zone was defined as the area where the depth is bellow or equal to 2 m. To parameterise the mass balance model (equation 3), a non-linear Bayesian model with informative priors was used. The parameters estimated were *I_0_* (eDNA generation coefficient), *b* (allometric scaling coefficient), and *k* (a combined degradation and sedimentation constant). Although equation 3 has both the degradation constant *k_deg_* and the sedimentation constant *k_sed_*, the current state of knowledge on eDNA degradation and sedimentation does not allow for a precise estimate of the values for *k_deg_* and *k_sed_*, and so one “disappearance constant” *k* was parameterised in the model as the sum of both (equation 4).

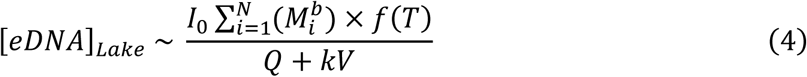

The model was fitted with a Gaussian distribution, four chains, 10 000 iterations, and *adapt_delta = 0.99*. A lake specific and temperature dependant *k* parameter was estimated to account for potential differences in degradation rates between lakes and in time, and a sampling period specific eDNA generation coefficient (*I_0_*) to account for season and sampling period effect on mean Brook Trout eDNA concentrations observed across lakes (Gaudet-Boulay et al., 2023), either due to ecological differences in the eDNA production coefficient (I_0_) across seasons or potential (unintentional) methodological differences in sampling. For all analysis, 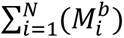 was approximated using the Taylor expansion for the first moment (Beaulieu et al., 2024) such that equation 4 becomes:

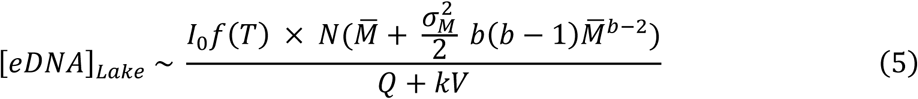

Where 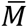 is the mean mass of the fish and 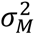 is the variance of the mass of the fish. The priors used were:

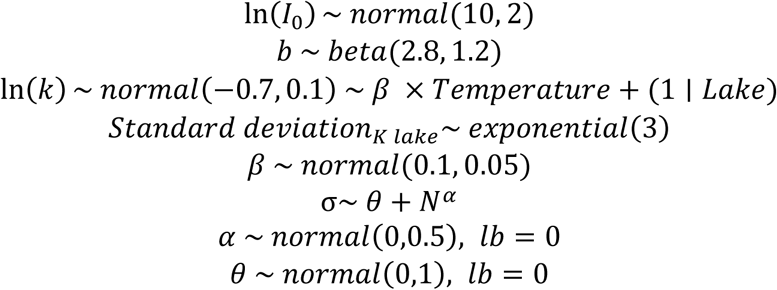

The prior on *I_0_* ensured posterior predictions were the correct order of magnitude, the prior on *b* ensured values were between 0 and 1 with a higher probability density around 0.7, which is approximately the *b* value estimated from Brook Trout bioenergetic models and expected based on bioenergetic theory (Deslauriers et al., 2017). The prior for *k* ensured values corresponding roughly to the expected values based on Lamb et al. (2022). eDNA data were scaled to Kilocopies m^-3^ to avoid working with very large numbers. The prior on sigma accounted for heteroscedasticity and increases variance with total abundance (N), and *α* and θ have a lower bound (*lb*) at zero.

### Comparing the mass balance eDNA approach to other existing models

Most previous studies examining relationships between eDNA concentrations and organism abundance/biomass typically directly regress observed eDNA concentrations against estimates of abundance and/or biomass (Yates, Glaser, et al., 2021). To evaluate the relative importance of accounting for both bioenergetics and particle dynamics (i.e., mass balance modelling), we also fitted models in which eDNA concentrations were directly related to both population biomass (equation 6) and allometrically scaled biomass (equation 7 and Yates, Cristescu, et al., 2021; Yates, Glaser, et al., 2021). Both models were fitted assuming a Gaussian distribution:

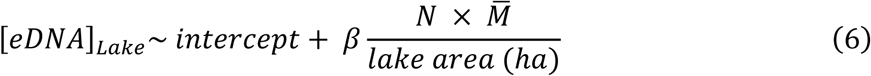

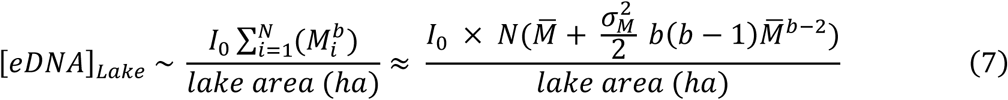

With the priors:

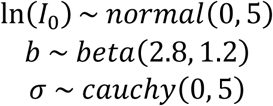

The performance of the different models was compared using the Bayes R^2^, looIC, and the value of the slope comparing the predicted and measured eDNA concentrations (the closer to one the better).

### Evaluating the efficacy of eDNA to estimate population abundance

We used our integrated bioenergetics/mass-balance approach to derive estimates of population abundance to compare their precision and accuracy relative to estimates of abundance derived from mark-recapture. Specifically, we were interested in evaluating the efficacy of eDNA to both i) differentiate total abundance *across* study populations; and ii) detect changes in total abundance *within* a population over time (i.e., population reductions due to harvesting). To produce posterior estimates of total abundance for each of the populations during each sampling period using the mass balance approach, *N* was isolated from equation 5 and the model was refitted with the same priors. Additionally, season and sampling period can have a strong effect on mean Brook Trout eDNA concentrations observed across lakes (Gaudet-Boulay et al., 2023), either due to ecological differences in the eDNA production coefficient (I_0_) across seasons or potential (unintentional) methodological differences in sampling. A sampling-period specific I_0_ was therefore also fitted in the model. Equation 5 then becomes:

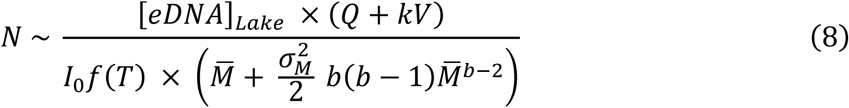

With the priors:

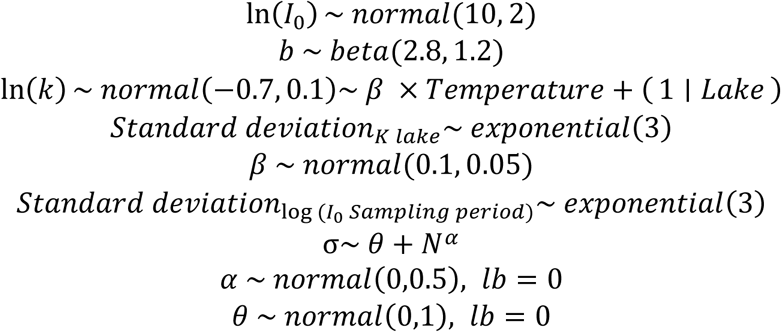

In this model, we also accounted for the measurement error on the mark-recapture total abundance estimates (McElreath, 2018). We also examined the efficacy of the integrated eDNA model to differentiate population biomass, which can be estimated from the integrated eDNA model (equation 8) for each population by multiplying the posterior distribution of *N* values by the mean mass of each population. Note that these models do not test the relationship between eDNA concentration and numerical abundance, but instead our ability to estimate numerical abundance from the markers’ concentration when considering a known mass distribution. The performance of the model was assessed using Bayesian R^2^ and the number of abundance estimates from eDNA that overlapped with mark-recapture (and their respective 95% credible intervals).

### Examining potential sources of bias

To assess the presence of systematic bias of the model, a Bayesian mixed model with non-informative priors was fitted with the z-score of each eDNA prediction as the independent variable. The fixed effects were mark-recapture abundance estimates, elevation, lake thermal mixing status (yes = stratified or no = mixed), season (Summer or Fall), lake area and experimental treatment (harvested or reference lake). We included interactions between thermal mixing status and elevation, between season and treatment, and between season and elevation. Lake was included as a random intercept to account for non-independence. This bias testing model was performed on the outputs of the eDNA mass-balance models, including sampling period specific *I_0_*.

### Can the nine lakes calibrate a model to predict abundance in novel systems?

Due to repeated sampling within and across years, the model described in equation 8 can estimate lake-specific degradation constants and sampling period specific *I*_0_, and is thus well-suited to monitoring abundance within and across this specific set of nine study lakes. However, this model cannot be extended to estimate population abundance in a lake without pre-existing data (e.g., where a lake specific *k* is unknown). To explore the efficacy of this approach to estimate abundance in lakes (equation 8) without pre-existing baseline data, we fitted two additional models: i) a model with lake specific k but no sampling period specific *I*_0_ to assess the importance of accounting for local, lake-specific eDNA degradation rates (referred to as equation 8.1 in the results) and ii) a model without a lake specific eDNA disappearance constant (*k*) nor sampling period specific *I*_0_ to assess how a generalizable model would perform as well as the importance of a sampling period specific eDNA generation coefficient (referred to as equation 8.2 in results). The performances of these different models were compared using Bayesian R^2^ values. All analyses were performed in *R* (CoreTeam, 2020) using the *brms* package (version 2.18.0) (Bürkner, 2018).

## Results

Brook Trout abundance varied significantly between lakes with estimates ranging from 72 (46 – 117) to 3173 (2644 – 3965) individuals based on mark-recapture. Temporal variation in population abundance was particularly pronounced in the size-selectively harvested lakes, where annual harvest ranged from 46.8% to 92.7% of the population. Despite these high harvest levels, compensatory effects allowed populations to partially recover between harvesting treatments (Matte et al., 2023). Natural mortality also contributed to population changes. For instance, in Dog Lake, population abundance declined by 32% between the Summers of 2018 and 2019, reflecting substantial non-harvest losses.

### Comparing the mass balance eDNA approach to other existing models

The eDNA mass balance model explained more variability in eDNA concentration and was more accurate than a linear regression with biomass or allometrically scaled biomass (Table 1, Fig. 2). While all three models had little bias, biomass alone explained only 24% of the variation in predicted eDNA values (Bayes R^2^). This increased to 44% after integrating the effect of allometry on eDNA production (e.g., allometrically scaled mass). Integrating both bioenergetics and mass-balance modelling approaches substantially improved upon this, with the integrated model explaining 71% of the variation in observed eDNA concentrations.

**Figure 2.**
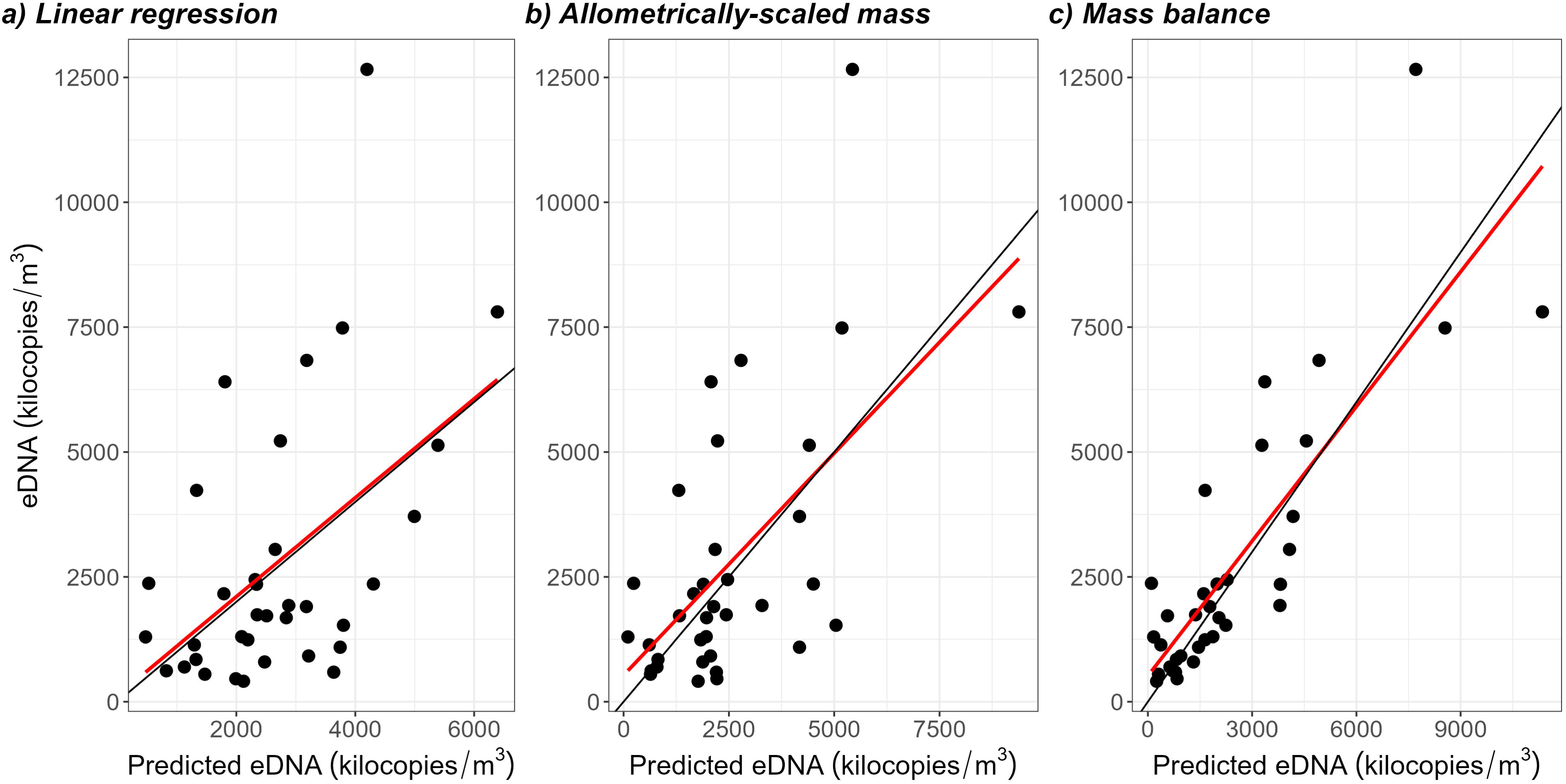
Comparison of model performance relating eDNA concentration to total population abundance. The black line represents the 1:1 slope, while the red line shows the linear regression between the predicted and measured eDNA concentrations. Parameter estimates and R^2^ values for each model are provided in Table 2.

**Table 1.** Parameter estimates and Bayes R^2^ for the linear regression, allometrically scaled biomass regression and the mass balance model. The formula column refers to the formula of the model specified as equations in the method section. The numbers in parentheses represent the 95 % credible interval on parameter estimates. The Bayes R^2^ reported are the mean of the Bayes R^2^ of each model iteration. *b* is the allometric coefficient, *k* is the sum of the degradation and sedimentation constant, *I_0_* is the eDNA generation coefficient, *β* is the slope of the linear model, sigma is the variance. The coefficient *β_ln(k) Temperature_* is the regression coefficient of the effect of water temperature on *k*. Sd stands for standard deviation.

| Model | Bayes $R^2$ | looIC | $b$ | $\ln(k)$ | Sd($\ln(k)$) | $\beta_{\ln(k) \text{ Temperature}}$ | $\ln(I_0)$ or $\beta$ | Sd($\ln(I_0)$) | Sigma | Formula |
| --- | --- | --- | --- | --- | --- | --- | --- | --- | --- | --- |
| Linear regression | 0.24 | 650.4 | - | - | - | - | 0.13 (0.05-0.21) | - | 2387 (1889-3037) | equation 6 |
| Allometrically scaled biomass | 0.44 | 645.0 | 0.58 (0.25-0.89) | - | - | - | 0.03 (-1.4-1.43) | - | 2162.93 (1707-2775) | equation 7 |
| Mass balance eDNA | 0.71 | 640.5 | 0.67 (0.28-0.98) | -0.70 (-0.90-0.51) | 0.29 (0.02-0.70) | 0.1 (0.02-0.18) | 10.52 (8.94-12.31) | - | $\theta = 5.13$ (4.46-5.88)<br>$\alpha = 0.13$ (0.06-0.17) | equation 5 |
| Mass balance with N as dependent variable, with lake specific and temperature dependant $k$ and sampling period specific $I_0$ | 0.75 | 550.8 | 0.79 (0.45-0.98) | -0.76 (-1.74-0.21) | 0.30 (0.06-0.65) | 0.12 (0.06-0.18) | 10.36 (8.85-12.11) | 0.19 (0.01-0.60) | $\theta = 1.94$ (0.52-3.35)<br>$\alpha = 0.18$ (0.13-0.22) | equation 8 |
| Mass balance with N as dependent variable without sampling period specific $I_0$ | 0.75 | 547.0 | 0.8 (0.49-0.98) | -0.7 (-0.9--0.51) | 0.3 (0.06-0.66) | 0.13 (0.07-0.19) | 10.46 (9.28-12.01) | - | $\theta = 2.14$ (0.83-3.47)<br>$\alpha = 0.18$ (0.12-0.22) | equation 8.1 |
| Mass balance with N as dependent variable without lake specific $k$ nor sampling period specific $I_0$ | 0.51 | 563.6 | 0.47 (0.06-0.96) | 0.49 (0.4-0.59) | - | - | 10.60 (8.23-12.47) | - | $\theta = 2.75$ (1.55-3.98)<br>$\alpha = 0.17$ (0.11-0.21) | equation 8.2 |

**Table 2.** Parameter estimates for bias testing. The Bayes R^2^ reported is the mean of the Bayes R^2^ of each draw. The dependent variable is the z-score of each total Brook Trout population abundance estimate, calculated as the estimate from eDNA minus the estimate from mark-recapture divided by the standard deviation of the draws of the estimate from eDNA. CI stands for credible interval.

| Variable | Parameter estimate | CI (95 %) | Bayes $R^2$ |
| --- | --- | --- | --- |
| Total abundance | -0.00018 | -0.0017 – 0.0013 | 0.33 |
| Lake area | -0.026 | -0.22 – 0.22 |  |
| Mixing (yes) | 0.075 | -1.89 – 2.03 |  |
| Elevation | -0.00043 | -0.0032 – 0.0024 |  |
| Season (Fall) | -0.35 | -2.27 – 1.57 |  |
| Treatment (Harvest) | 0.3 | -1.16 – 1.76 |  |
| Elevation*Mixing | -0.0008 | -0.0023 – 0.0008 |  |
| Elevation*Season | 0.0007 | -0.0008 – 0.0024 |  |
| Season*Treatment | 0.87 | -0.77 – 2.51 |  |

### Evaluating the efficacy of eDNA to estimate population abundance

Total abundance estimates derived from eDNA concentration data (equation 8) effectively differentiated populations across spatial and temporal scales (Fig. 3). The study populations exhibited a substantial gradient in total abundance across lakes, with the smallest population exhibiting a 32-fold difference in abundance relative to the largest population. At this scale, the integrated eDNA model was very effective at differentiating abundance across populations (Fig 3).

**Figure 3.**
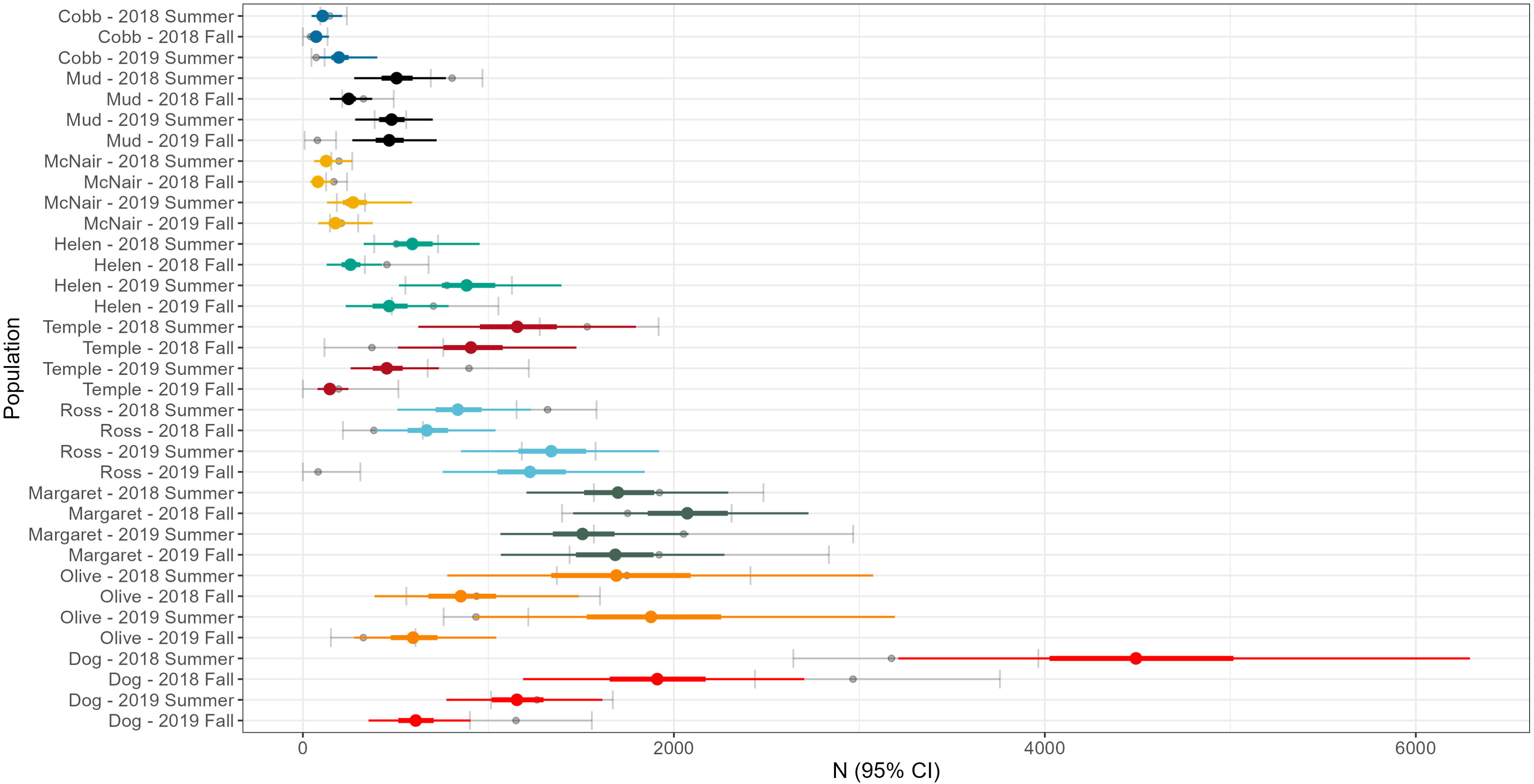
Total population abundance estimates from eDNA for all the populations and sampling periods. The thick colored line represents the 50 % credible interval of eDNA estimates, while the thin colored lines represent the 95 % credible intervals. Grey lines and dots represent the 95% credible interval and mean from the mark-recapture abundance estimates. eDNA estimates are derived from the full mass balance model (Equation 8).

Variations in abundance were significantly greater among populations than within populations, even for populations that exhibited substantial within-population declines due to size selective harvesting manipulations. Within a population, abundance levels were more consistent across seasons and years (Fig. 3 and Fig. 4). Yet, even within populations, the accuracy and precision of the integrated eDNA model was sufficient to detect changes across seasons and years in several of the harvested lakes (e.g., Olive and Temple), as well as natural fluctuations in abundance across control populations (e.g., Margaret and Dog lakes).

**Figure 4.**
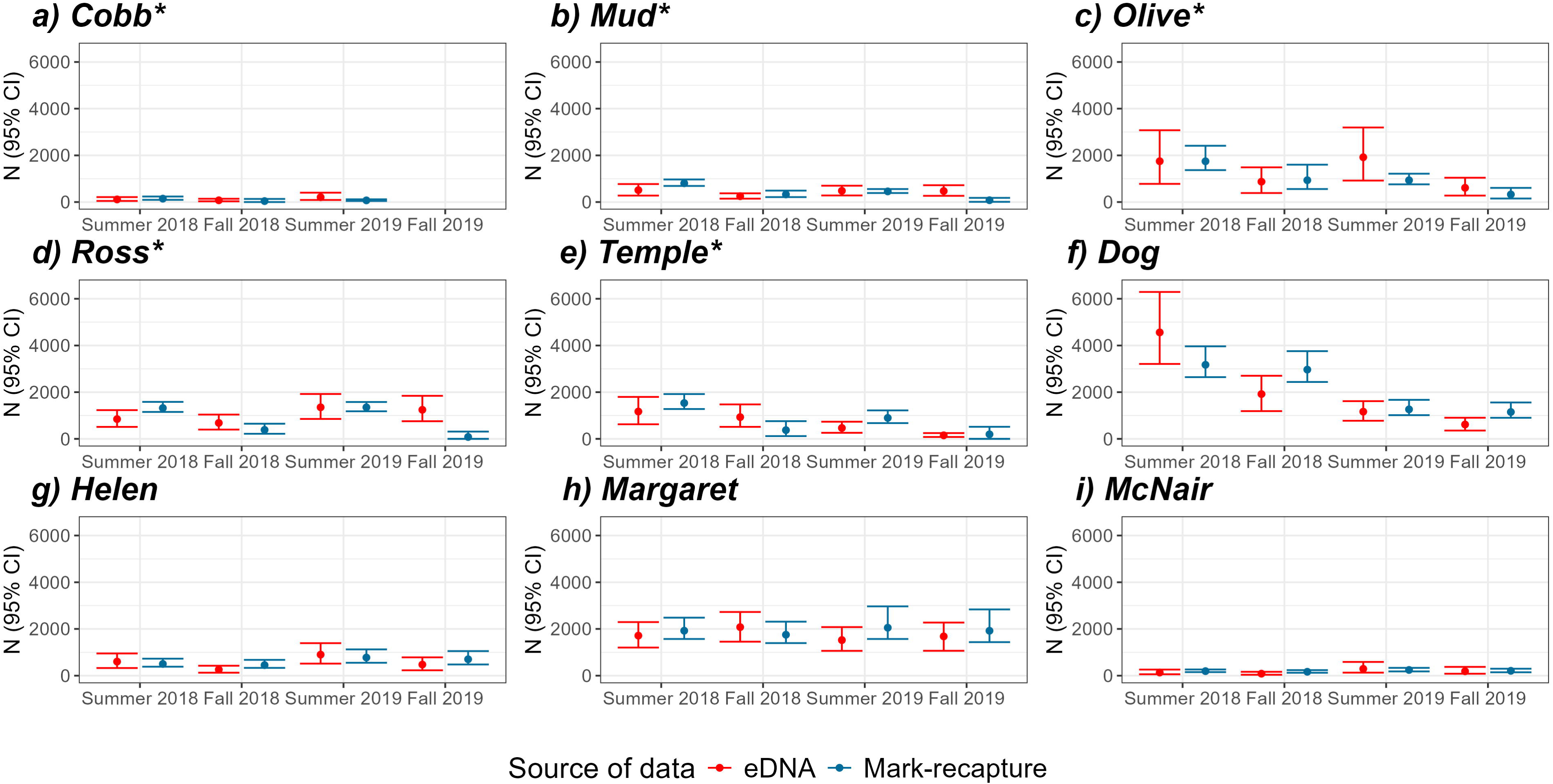
Total population abundance estimates from the eDNA mass balance model (red) and mark-recapture (blue). eDNA estimates are produced from the fitted mass balance model with sampling period specific eDNA generation coefficient (I_0_) (Equation 8). Lakes that were harvested are marked with an asterisk (*).

The majority (32/35 or 94%) of the total abundance estimates derived from the eDNA mass balance model (equation 8) fell within the 95% credible intervals for total abundance estimated by mark-recapture (Fig. 3 and 4). The integrated eDNA model similarly proved effective at differentiating population biomass across space and time, also overlapping with mark-recapture estimates in 32/35 of the cases. The biomass estimates from the integrated eDNA model were highly correlated with estimates derived from mark-recapture (Bayes R^2^ = 0.89) (Fig. 5) and, as with abundance, enabled the clear differentiation of biomass both across and within populations (Fig. SM1).

**Figure 5.**
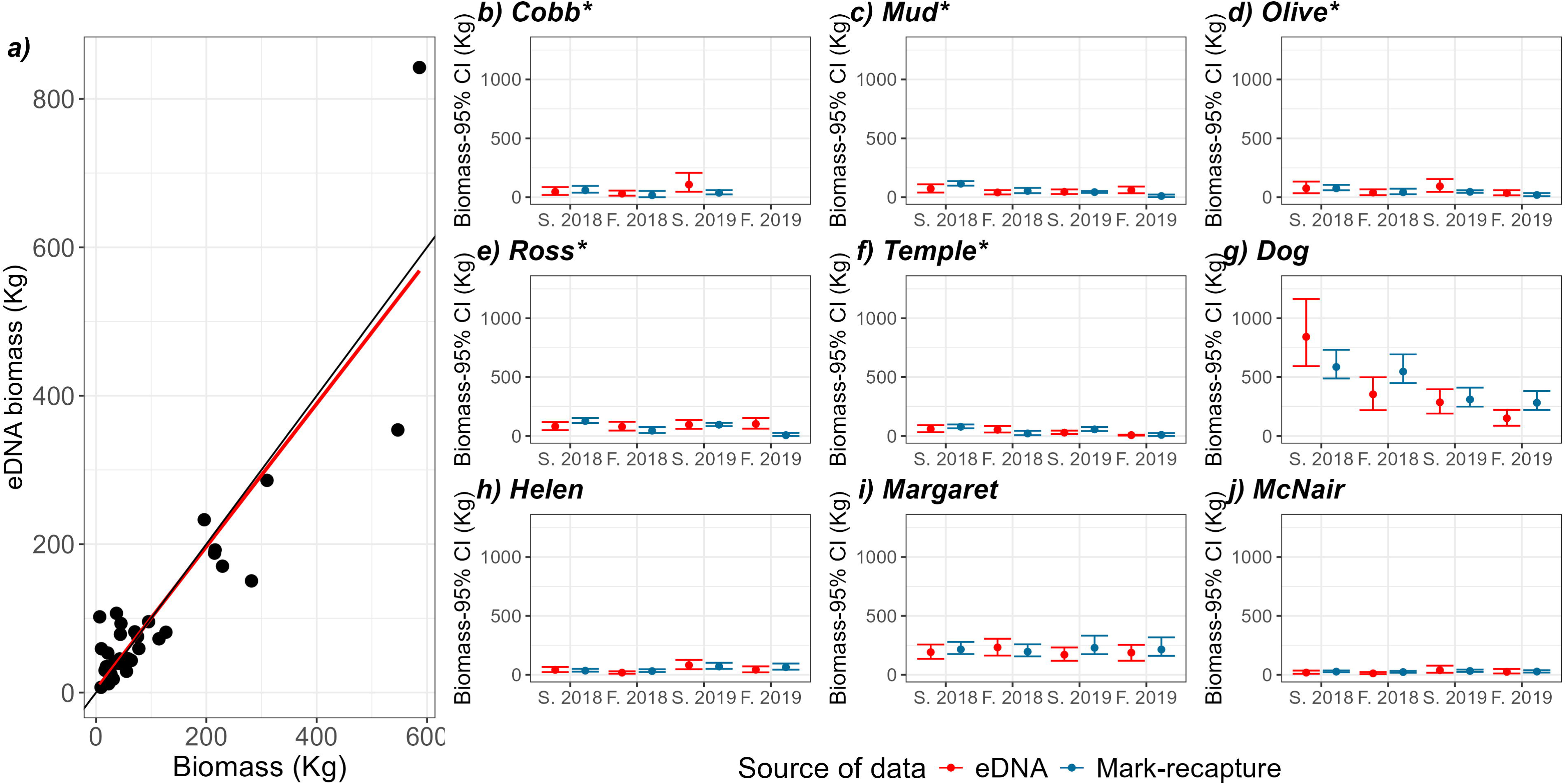
Population biomass estimated from the eDNA mass balance model and mark-recapture. In a) the x axis is the biomass estimated from the mark-recapture abundance estimates and the y axis is the biomass estimated from the eDNA mass balance model. The red line is the linear regression between the two, and the black line is the 1:1 slope. In b) to j) the biomass estimated from the eDNA mass balance model (red) and the biomass estimated from the mark-recapture (blue). eDNA estimates are produced from the fitted mass balance model with sampling period specific eDNA generation coefficient (I0) (equation 8.2). Lakes that were harvested are marked with an asterisk (*).

### Examining potential sources of bias

When testing for different biases of the eDNA total abundance estimates, there was no significant systematic bias detected related to the total Brook Trout abundance nor lake size. There was a non-significant tendency to generate larger abundance estimates in lakes experiencing experimental harvesting during the Fall seasons (e.g., Ross and Mud at Fall 2019) compared to mark-recapture (Fig 4, Table 2, and Appendix 5).

### Can the nine lakes calibrate a model to predict abundance in novel systems?

Incorporating a lake specific eDNA disappearance constant (*k*; the sum of the sedimentation and degradation constant) significantly improved model performance, as demonstrated by the large standard deviation on a general *k* estimated across lakes and the reduction in model performance when removing lake specific *k* (equation 8.2), with the Bayes R^2^ dropping from 0.75 to 0.51 (Table 1 and Appendix 6). The effect of accounting for a sampling period specific eDNA generation coefficient (I_0_; equation 8.1) had no effect on model performance. Brook Trout abundances predicted from the eDNA mass balance model that lacked a sampling period specific I_0_ (equation 8.1) also fell within the 95% credible intervals for total abundance estimated by mark-recapture for 31/35 estimates (Appendix 5).

## Discussion

There is an increasing demand for abundance data to understand and manage ecosystems (Huang et al., 2022). Yet, the application of eDNA to aid in quantifying the abundance or biomass of aquatic organisms remains a controversial topic (Parsley et al., 2024; Rourke et al., 2022; Yates et al., 2019). This stems from the complex interplay of biotic and abiotic factors that influence eDNA production, degradation, and transportation in natural aquatic environments (Yates, Cristescu, et al., 2021). These are considerably more variable across space and time in natural ecosystems relative to controlled laboratory settings (Yates et al., 2019).

Herein, we demonstrate that eDNA can provide estimates of organism abundance and biomass comparable in precision and accuracy to mark-recapture. However, achieving this accuracy required accounting for biotic and abiotic factors that affect its distribution in natural ecosystems, namely the ecology of its production and the dynamics of the physical eDNA particles. Applying our integrated bioenergetics and mass balance approach to model quantitative eDNA signals greatly improved the relationship between eDNA concentration and total abundance estimates in the study ecosystems, compared to a basic linear correlation between either eDNA and biomass or eDNA and allometrically scaled biomass. Our integrated approach allowed for the accurate differentiation of population abundance and biomass across study populations (i.e., space) based on lake eDNA concentrations. Quantifying the concentration of eDNA within the study lakes also facilitated the prediction or monitoring of temporal changes in fish abundance/biomass over time within several populations.

Previous research has demonstrated the influence of several factors on eDNA concentration and dynamics, including transportation (Carraro et al., 2018; Sepulveda et al., 2020; Shogren et al., 2017), degradation (Carraro et al., 2018; T. Jo et al., 2019; T. Jo & Minamoto, 2021; Strickler et al., 2015), temperature (Caza-Allard et al., 2022; T. S. Jo, 2023), and fish metabolic rate (Yates, Cristescu, et al., 2021; Yates et al., 2023; Yates, Wilcox, et al., 2021). However, our study is the first to integrate mass balance processes with bioenergetics to relate eDNA concentration to total population abundance in natural ecosystems. We empirically demonstrate that eDNA could represent a time- and cost-effective tool for generating total abundance estimates, with sufficient precision to detect changes in abundance, by integrating relatively simple-to-collect environmental and ecological data.

### Evaluating the efficacy of eDNA to estimate population abundance

eDNA abundance estimates corresponded to mark-recapture abundance estimates in 32 out of 35 cases (91%) when considering the 95% credible interval. All three discrepancies were likely due to issues associated with our unique study design causing the violation of assumptions in our mass-balance modelling (i.e. experimental harvesting), and/or the violation of assumptions associated with the mark-recapture estimates for populations in the Fall sampling periods (e.g., no natural mortality or emigration).

There was a non-significant trend wherein estimates of abundance derived from eDNA tended to be higher than for mark-recapture in harvested lakes in the Fall. The time necessary for eDNA concentrations to stabilize depends on eDNA degradation rate, which can be impacted by amplicon size, temperature, and potentially UV radiation, pH, and microbial activity (Lamb et al., 2022). The values estimated by the eDNA mass balance model for the *k* degradation parameter (the eDNA “disappearance rate”, or the sum of the sedimentation and degradation rates) corresponded to expectations based on previous controlled experiments. Notably, meta-analysis reported degradation constant values between 0–0.4 h^-1^ or 0 – 9.6 day^-1^ in freshwater ecosystems (T. Jo & Minamoto, 2021; Lamb et al., 2022). We obtained an average *k* value estimate ranging from 1.3 to 6.9 day^-1^ depending on lakes and water temperature, with a mean of 3.06 day^-1^. Based on the eDNA disappearance constant *k* estimated by the model, the outflow rate, and the Summer eDNA concentration in each lake and each year, we estimated that one to fifteen days would be necessary for eDNA turnover in this study system (Appendix 8). For logistical reasons (i.e., the harvesting treatment took a substantial amount of time and extended late in the season), there were generally fewer than seven days between the end of harvesting and eDNA sampling within several of the harvested lakes, which is less than the turnover rate in most cases. This was especially the case for Mud and Ross lakes in Fall 2019, which represent two of the three significant discrepancies between eDNA and mark-recapture. A tendency for the integrated eDNA model to overestimate fish abundance following harvesting could be explained by many eDNA copies being released during harvesting. Carcasses can be a significant source of eDNA (Dunker et al., 2016) and often remained in nets for 24 - 48h, thus increasing the initial amount of eDNA and time before stabilisation back to a steady state concentration.

There was also one case where the eDNA estimate weas lower than mark-recapture estimates in Dog Lake in Fall 2019. An instinctive response may be to attribute this discrepancy to error in the integrated eDNA model. However, this discrepancy between the eDNA-derived and mark-recapture abundance estimates could be attributable to violations of the assumptions of the mark-recapture methodology employed. Mark-recapture estimates demonstrated that Dog Lake exhibited a substantial natural decline in abundance between 2018 and 2019. While mortality was assumed to be negligible between Summer and Fall, we do not actually know when this mortality event occurred because mark-recapture studies occurred in the spring and Summer of each year. Results from the eDNA model could imply that the natural mortality event may have occurred during both Summer 2018 and 2019. In these cases, eDNA may more accurately reflect the abundance of fish in the study lakes than mark recapture estimates, for which we assumed there was no natural mortality nor emigration for the Fall estimates.

### Generalisation of the integrated bioenergetic and mass balance model

Although there were a few discrepancies between eDNA and mark-recapture derived estimates in abundance, these discrepancies still reflected a relatively high level of precision *across* study populations, for which eDNA permitted broad differentiation across high- and low-abundance populations. Although we advise caution, these results imply that eDNA can be used to reliably estimate abundance and biomass in nature across populations. Furthermore, mark-recapture estimates themselves can exhibit biases or fail when assumptions are violated. Our study design does not allow us to identify which estimate (eDNA vs. Mark-recapture) is closer to the ‘true’ population size, and these issues would only be compounded in larger natural ecosystems. Overall, we detected no systematic bias in eDNA-derived estimates related to lake size, mixing, elevation, and population size. The integrated bioenergetics and mass balance approach that we applied to the analysis of eDNA concentrations produced mostly accurate population abundance estimates in small to medium-sized oligotrophic lakes. With further validation, such models have the potential to be extended to other aquatic ecosystems.

Accounting for lake mixing was an important component of the mass-balance modelling because the volume of the waterbody is a critical parameter in mass-balance modelling approaches. Previous research has demonstrated that eDNA disperses throughout a lake and is roughly homogenous during thermal mixing (Littlefair, Hrenchuk, et al., 2021). During lake stratification, we found that the volume of the epilimnion is the appropriate volume input data, likely because Brook Trout remains close to the surface during Summer (Tiberti et al., 2017). In contrast, when a lake is undergoing thermal mixing, the volume of the entire lake is the appropriate volume input data in the mass-balance model as eDNA is diluted across the entire lake, rather than just the thermal layer inhabited by the target species (Littlefair, Hrenchuk, et al., 2021). We detected no systematic bias related to lake thermal mixing and mixing explained no discrepancies between eDNA and mark-recapture abundance estimates. The absence of overestimation of Brook Trout abundance during mixing implies that the resuspension of sedimentary DNA during lake mixing likely had a negligible impact on abundance estimates. This is despite lake sediments represent a potentially potent source of eDNA (Turner et al., 2015). However, the amount of eDNA stored within sediments and how it exchanges with the water column also depends on an organism’s spatial distribution, as well as biotic and abiotic conditions in water and at the water-sediment interface (Capo et al., 2021). Sedimentary DNA re-suspension may be more important in other types of lakes or for other species, particularly for systems inhabited by benthivorous fish species that disturb sediment during feeding (e.g. Parkos et al., 2003).

### Future directions: Can the nine lakes calibrate a model to predict abundance in novel systems?

Our perspective on the future application of eDNA to estimate population abundance and biomass is not to replace conventional methods, but rather to complement and/or improve upon estimates generated by them. Notably, the integrated bioenergetic and mass-balance model developed herein requires population size structure data which can only be obtained using mixed mesh index gillnetting. These methods, often used in state and provincial monitoring programs to estimate relative abundance based on CPUE (e.g. Zimmerman & Palo, 2011), require fewer resources than mark-recapture techniques. To broadly apply eDNA to estimate abundance using the methods and models developed herein also requires the establishment of a ‘calibration system’, in which abundance estimated using conventional methods on a subset of ecosystems are used to calibrate an eDNA/abundance model. Once calibrated, this model can be applied to estimate abundance in other systems or time points lacking conventional estimates, by sampling eDNA and the required physicochemical parameters.

However, our findings imply that the capacity to identify local predictors of *k* would be particularly critical for such applications, as baseline data to estimate *k* rates within an individual system substantially improved our model performance. To increase the reliability of abundance estimates in freshwater systems derived from eDNA, further research is needed on eDNA dynamics in lentic systems, with a particular focus on drivers of sedimentation and degradation rates. A quantitative understanding of these interactions would allow us to better predict sedimentation and degradation rates specific to local and temporal conditions. This would in turn likely improve the integrated eDNA bioenergetic and mass-balance model performance as such dynamics could be integrated into the modelling process. Additionally, such knowledge would facilitate the estimation of the disappearance *k* parameter for lakes and time points outside of any putative “calibration system” lakes simply by measuring the relevant environmental variables. Alternatively, a lake-side small scale degradation experiment to estimate *k* could reduce the amount of mark-recapture data needed to calibrate the model as the physiological parameters are more stable in space and time.

A growing number of individual studies on eDNA degradation have found that eDNA degradation is the result of a complex combination of many variables such as pH, temperature, UV radiation and microbial activity, and eDNA properties such as amplicon size and whether the targeted gene is nuclear or mitochondrial (T. Jo et al., 2019; T. Jo & Minamoto, 2021; Saito & Doi, 2021b, 2021a; Strickler et al., 2015). Although temperature has emerged as a consistently impactful parameter affecting eDNA degradation rates, the importance of other parameters such as UV, microbial activity, eDNA source, and amplicon size remains to be confirmed (T. Jo & Minamoto, 2021; Lamb et al., 2022; Saito & Doi, 2021b). Collectively, we are still far from producing effective eDNA degradation models, as few studies to date have examined the interaction between these different environmental variables and eDNA properties (especially in natural ecosystems) and even fewer have examined eDNA sedimentation. However, in the absence of controlled empirical studies, further large-scale observational studies in natural ecosystems could facilitate the identification of drivers of local eDNA decay rates. Experiments studying degradation dynamics in natural ecosystems are thus crucial to facilitate and improve the adoption of eDNA to estimate abundance in natural ecosystems.

Theoretical and empirical evidence suggests that eDNA production rate is related to fish metabolic rate (Hervé et al., 2023; Thalinger et al., 2021; Yates et al., 2023; Yates, Glaser, et al., 2021). However, to our knowledge, only one study to date has experimentally verified this relationship under controlled empirical conditions and found a positive correlation between fish activity rate, energy use, and eDNA shedding rate (Thalinger et al., 2021). The mass balance model assumes eDNA production scales with body mass and is influenced by temperature-dependent metabolic rates in poikilothermic Brook Trout (Hervé et al., 2023; Thalinger et al., 2021; Yates, Cristescu, et al., 2021; Yates et al., 2023). We used a temperature-dependent metabolic function (Hartman & Cox, 2008) to adjust for temperature effects on eDNA, though this may not fully reflect eDNA production dynamics. While many studies have documented an effect of temperature on eDNA production (e.g. Caza-Allard et al., 2022; T. Jo et al., 2019), whether this effect directly reflects metabolic responses to temperature remains to be tested. Directly quantifying the eDNA production temperature dependency function could thus improve model assumptions and performance.

Our study additionally included mostly headwater lakes, so eDNA inputs from upstream were considered equal to zero. However, this is rarely the case within a watershed. To add eDNA transportation into the lake within the integrated model, data on both the inflow rate and the average eDNA concentration in the inflow are required. In theory, these data can be implemented in equation 4 (Chapra & Reckhow, 1983) such that:

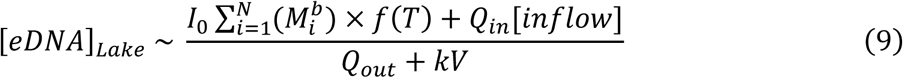

Where *Q*_*in*_ is the inflow rate, *Q*_*out*_ is the outflow rate, and [*inflow*] is the mean eDNA concentration of the inflow.

## Conclusion

Our study proposes an integrated bioenergetics and mass-balance approach to relate eDNA concentration to population abundance estimation. Our integrated eDNA model performed well at estimating abundance, and it facilitated total abundance and biomass estimates that were comparable to conventional mark-recapture methodologies, often considered a ‘gold-standard’ for population abundance estimates. We show that eDNA can detect both spatial and temporal differences in fish abundance, provided the equilibrium assumptions of the model are respected. To respect these assumptions, we recommend avoiding sampling ten to twenty days after known fluctuations in abundance (e.g. harvest), depending on the eDNA degradation rate and outflow flux. Overall, the results of this study represent an important advancement in the use of eDNA to quantify total population abundance, with strong implications for the application of eDNA for abundance estimates in a context of ecosystem management. The quantitative analysis of eDNA, applied with an integrated bioenergetics and mass-balance approach, could provide a cost-effective, precise, and accurate tool for managers and researchers. This is critical in an era where the acquisition of population abundance data is increasingly important for biodiversity management (e.g. Canals et al., 2021; Fonseca et al., 2023; Kelly et al., 2024).

## Supporting information

supplement_beaulieu-et-al-2024

## Acknowledgements

We thank Parks Canada for logistic support, Andrew Macdonald for statistical advice, Maria Gheta for the help with the Taylor expansions, Yves Prairie for math validation, Jérôme Lemelin for the map, and Mélanie Desrochers for GIS advice. We also express our gratitude to Louis Astorg, Dylan Glaser, Brent Brooks, Shannon Clarke, Jean-Michel Matte, Justin Budyk, Mélia Lagacé, Mathilde Salamon, Sydney Urschel, and Ahmad Yaghi for field work. Research funding was provided by a Fonds de Recherche du Québec-Nature et Technologie (FRQNT) team projet grant 254557 (AMD, MEC, and DJF), a grant from the Natural Sciences and Engineering Research Council of Canada (NSERC) Strategic Project grant STPGP 494015-16 (AMD, DJF), and a NSERC Discovery Grant 2022-03706 (AMD). We thank the Groupe de recherche interuniversitaire en limnologie (GRIL) and the Faculty of Science at the Université du Québec à Montréal for providing contributions to scholarship funding for JB. MCY was supported by a FRQNT postdoctoral fellowship.

## Authors contribution

JB designed the models and performed the analysis. JB and MCY contributed to the writing of the manuscript. MCY and JB collected the eDNA samples. MCY extracted the eDNA samples and conducted laboratory analyses. All authors contributed to the study design and subsequent edits to the manuscript.

## Data availability statement

Data and codes will be available upon manuscript acceptance. In the meantime they can be found at: https://share.multcloud.link/share/3380ba5d-8bda-40d8-a7c7-c60298e2f1d2

## Animal care

Fieldwork was conducted under a Parks Canada Agency Research and Collection Permit (KooNP-2017-24,999)

