## supplement_beaulieu-et-al-2024 for "eDNA provides accurate population abundance estimates with bioenergetics and particle mass-balance modelling"

PNAS

#### This PDF file includes:

Appendix 1: Studied lakes

Appendix 2: Contamination prevention

Appendix 3: Results when removing the two plates with low efficiency

Appendix 4: Temperature data used

Figure SM1: biomass estimates across lakes

Appendix 5: Results without the time period specific I0

Appendix 6: Population estimates when not including a K estimate for each lake nor sampling period specific I0.

Appendix 7: Bias testing with the mass balance model

Appendix 8: eDNA turnover time

### Appendix 1: Studied lakes

**Table S1**: Characteristics, locations, and experimental treatment category of the nine study lakes, ordered from highest elevation to lowest elevation. The pH and temperature data are from YSI® vertical profiles. Data from the first three meters for the whole study period were averaged to give an idea of the abiotic conditions.

| Lake | Latitude | Longitude | Elevation (m) | Treatment | Area (ha) | Max. depth (m) | pH | Temperature (°C) |
|---|---|---|---|---|---|---|---|---|
| Temple | 51.36604 | -116.17847 | 2207 | Harvest | 3.25 | 12.0 | 7.77 | 5.24 |
| Helen | 51.68436 | -116.41361 | 2400 | Reference | 2.48 | 15.0 | 7.95 | 8.60 |
| Margaret | 51.58135 | -116.37138 | 1808 | Reference | 18 | 28.2 | 8.19 | 6.44 |
| Ross | 51.43745 | -116.30307 | 1735 | Harvest | 6.61 | 21.5 | 8.08 | 6.69 |
| Mud | 51.44039 | -116.17628 | 1600 | Harvest | 7.2 | 7.2 | 8.31 | 14.0 |
| McNair | 51.40644 | -116.15767 | 1532 | Reference | 1.66 | 4 | 8.11 | 10.8 |
| Olive | 50.67336 | -115.93733 | 1470 | Harvest | 1.66 | 3.5 | 7.98 | 10.0 |
| Cobb | 50.66119 | -115.87817 | 1260 | Harvest | 2.29 | 8 | 8.13 | 15.9 |
| Dog | 50.78036 | -115.92953 | 1185 | Reference | 11.5 | 4.7 | 8.37 | 15.7 |

### Appendix 2: Contamination prevention

#### *Field and transport*

All lake samples were collected from an inflatable kayak that was decontaminated 48 h prior to sampling by a complete soaking in a 2% household bleach solution for 15 minutes. The kayak paddle and life jacket used were similarly decontaminated. All samples were collected with Whirl-Pak™ bags (Uline, Ontario, Canada) while the individual collecting the sample wore sterile nitrile gloves. The vacuum hand pump (Soil Moisture, California, USA) used for filtration was wiped with a 30% household bleach solution and allowed to rest for ten minutes before rinsing with distilled water before the sampling day. The filtering manifold components were soaked in a 30% household bleach solution for a duration of 8 minutes and rinsed with distilled water between each sample to avoid cross-contamination of eDNA. Filters were handled with a pair of metal forceps that were soaked in a 30% household bleach solution for 8 minutes between sample filtrations before rinsing in distilled water.

For transport, manifolds were transported in a backpack cooler (Polar Bear Coolers, Georgia, USA) whose interior was wiped with a 30% bleach solution and allowed to rest for ten minutes before rinsing with distilled water. Manifold components were stored in sealed individual plastic zippered bags for transportation to Hidden Lake and Corral Creek. Writing utensils (pencils and markers) were also wiped with a 30% bleach solution and stored in zippered bags. The cooler bag used to keep eDNA samples cold and transportation was decontaminated by wiping with a 30% household bleach solution and allowed to rest for 10 minutes before rinsing with distilled water. This freezer contained two frozen gel packs that were decontaminated by soaking in 30% household bleach solution for ten minutes and rinsing with distilled water. Filters were immediately transported to Kootenay crossing, where they were stored in a zippered plastic bag in a -20 ℃ freezer that was previously decontaminated by wiping with a 30% household bleach solution.

##### *Extraction*

To avoid cross-contamination between lake and stream sites, as well as time periods, extractions were conducted in batches corresponding to sampling period and lake or stream sites (i.e. only samples from the lake or only samples from streams from a single sampling period were extracted in a batch). Extracted DNA was eluted into 130 µL of AE buffer and stored in a clean -20 ℃ freezer solely dedicated to the storage of extracted eDNA product (i.e. no post-PCR products or tissue samples). All extractions were conducted in a room dedicated solely to the extraction of eDNA samples that is cleaned on a weekly basis with a 10% household bleach solution and is free from PCR products, animal tissues, or high-concentration DNA. All individuals entering and working in the room are required to wear dedicated and clean lab coats, dedicated footwear with shoe covers, hair nets, and nitrile gloves. Laboratory surfaces were soaked with a 20% household bleach solution for ten minutes before and after each extraction batch. PCR Clean Wipes™ (Thermo Scientific, Massachusetts, USA) were also used to decontaminate all lab surfaces and micropipettes before and after each extraction batch.

### Appendix 3: Results when removing the two plates with low efficiency

To make sure the two plates with low qPCR efficiency had no impact on the results, we fitted the mass balance integrated model without the data originating from these plates: data from Cobb Lake at Fall 2018, Temple Lake at Fall 2018, and McNair Lake in 2019. Removing these data had no significant impact on the performance of model, nor the population estimates (Fig. SM2 and SM3)

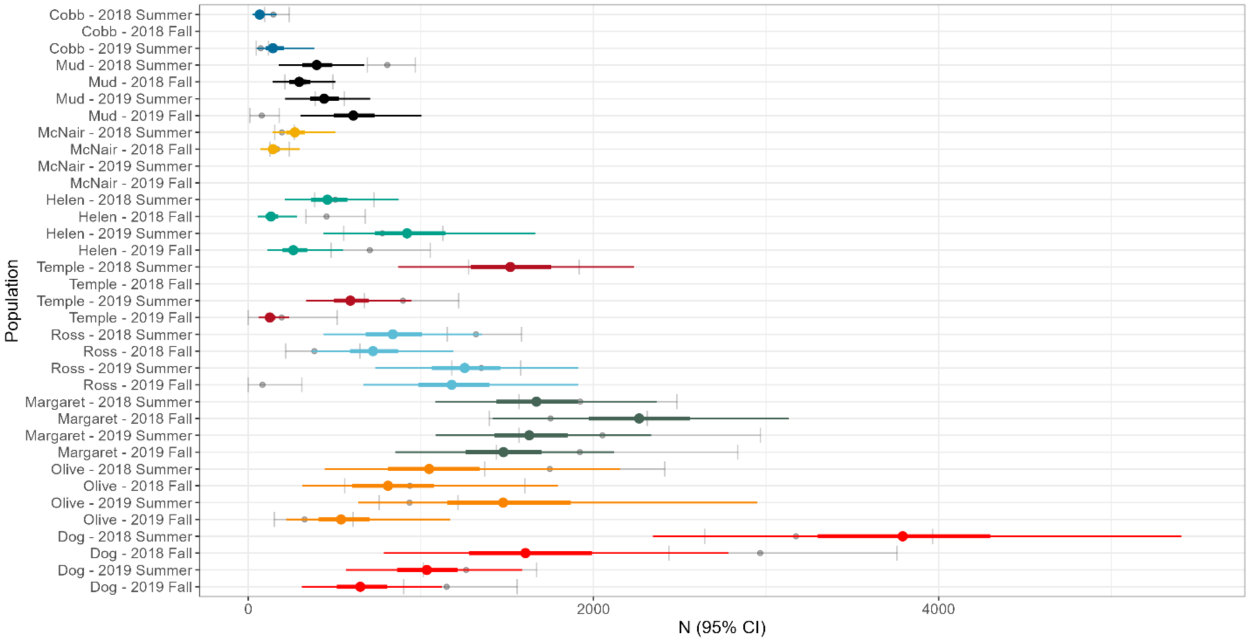

**Figure SM2**. Total abundance estimates from eDNA for all the populations and sampling periods. The colored thicker line represents the eDNA 50 % credible interval, and the colored thinner lines represent the eDNA 95 % credible intervals. The grey lines and dot represent the 95% credible interval and mean from the mark re-capture abundance estimates. eDNA estimates are produced from the full mass balance model (equation 8), but after removing the plates with low efficiency.

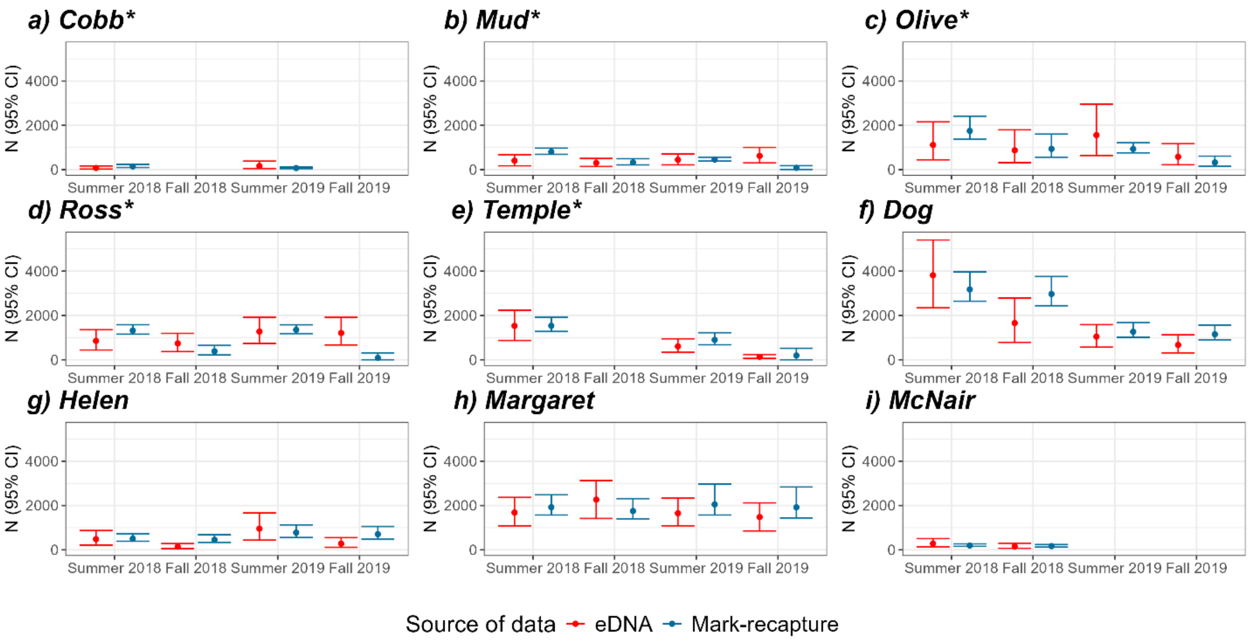


**Figure SM3.** Total abundance estimates from the eDNA mass balance model without sampling period specific *I0* (red) and mark-recapture (blue). eDNA estimates are produced from the fitted mass balance model (equation 8 in the main manuscript). The * mark lakes that were harvested. eDNA estimates are produced from the full mass balance model (equation 8).

### Appendix 4: Temperature data used

**Table SM1:** Source of the temperature data used for each lake and each time period.

| Lake | Time period | Data source | Days with datalogger | Average temperature (°C) |
|---|---|---|---|---|
| Temple | Summer 2018 | Datalogger | 14 | 7.04 |
| | Fall 2018 | Datalogger | 14 | 9.06 |
| | Summer 2019 | Datalogger | 18 | 7.30 |
| | Fall 2019 | 2018 Datalogger | 0 | 9.06 |
| Helen | Summer 2018 | Datalogger | 14 | 13.67 |
| | Fall 2018 | Datalogger | 1 | 14.65 |
| | Summer 2019 | Datalogger | 1 | 11.14 |
| | Fall 2019 | 2018 Datalogger | 0 | 14.65 |
| Margaret | Summer 2018 | Datalogger | 4 | 12.74 |
| | Fall 2018 | YSI | 0 | 9.30 |
| | Summer 2019 | Datalogger | 4 | 11.37 |
| | Fall 2019 | Datalogger | 7 | 11.59 |
| Ross | Summer 2018 | Datalogger | 14 | 12.14 |
| | Fall 2018 | Datalogger | 14 | 9.08 |
| | Summer 2019 | Datalogger | 27 | 10.98 |
| | Fall 2019 | 2018 Datalogger | 0 | 9.08 |
| Mud | Summer 2018 | Datalogger | 14 | 17.55 |
| | Fall 2018 | Datalogger | 14 | 12.19 |
| | Summer 2019 | Datalogger | 8 | 15.35 |
| | Fall 2019 | 2018 Datalogger | 0 | 12.19 |
| McNair | Summer 2018 | YSI | 0 | 10.78 |
| | Fall 2018 | YSI | 0 | 10.0 |
| | Summer 2019 | YSI | 0 | 9.25 |
| | Fall 2019 | 2018 YSI | 0 | 10.0 |
| Olive | Summer 2018 | Datalogger | 14 | 15.85 |
| | Fall 2018 | YSI | 0 | 9.88 |
| | Summer 2019 | Datalogger | 14 | 13.53 |
| | Fall 2019 | Datalogger | 13 | 11.39 |
| Cobb | Summer 2018 | Datalogger | 14 | 19.49 |
| | Fall 2018 | Datalogger | 14 | 15.79 |
| | Summer 2019 | Datalogger | 14 | 17.79 |
| Dog | Summer 2018 | Datalogger | 14 | 18.84 |
| | Fall 2018 | Datalogger | 14 | 15.64 |
| | Summer 2019 | Datalogger | 27 | 18.20 |
| | Fall 2019 | 2018 Datalogger | 0 | 15.64 |

### Figure SM1: biomass estimates across lakes

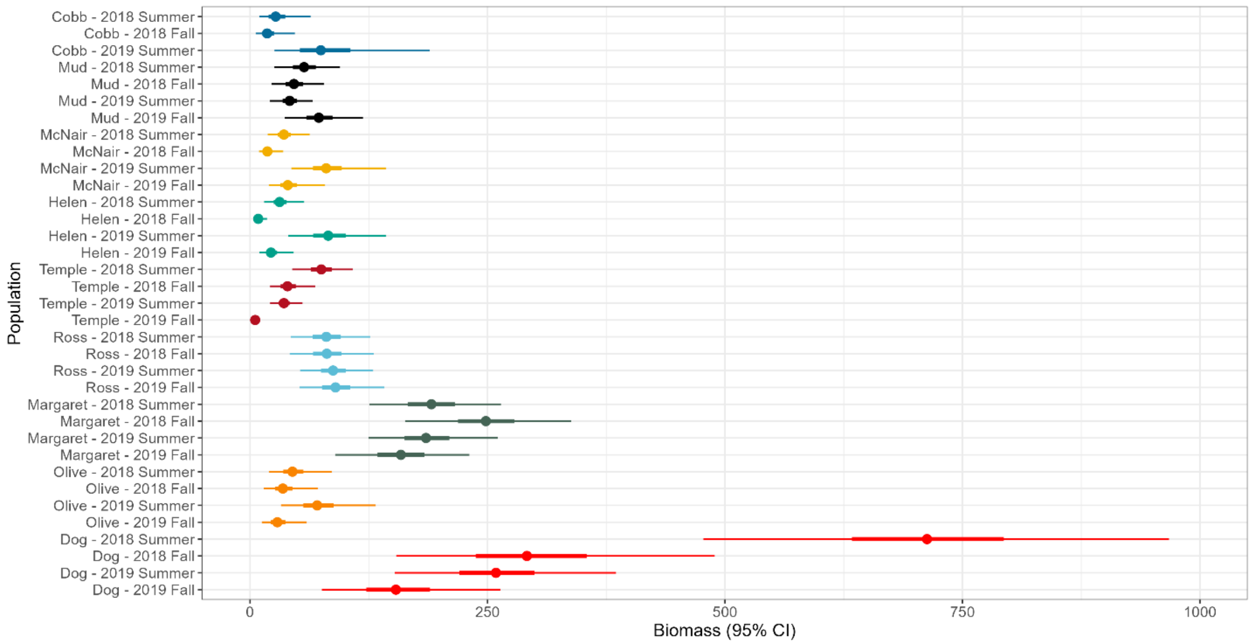


**Figure SM1**. Total abundance estimates from eDNA for all the populations and sampling periods. The thicker line represents the 50 % confidence interval, and the thinner lines represent the 95 % confidence intervals. eDNA estimates are produced from the full mass balance model (equation 8).

### Appendix 5: Results without the time period specific I0

To isolate and consider the changes in eDNA abundance estimates related to the summer 2018 sample thawing during transport, a sampling period specific eDNA generation coefficient *I0* was modeled in the results presented in the main manuscript. This annexe presents the results and figures with only one general *I0* estimated. Adding a sampling period specific I0 had very little impact on the general results but corrected slightly the estimates from Margaret Lake, Olive Lake and Ross Lake at fall 2018.

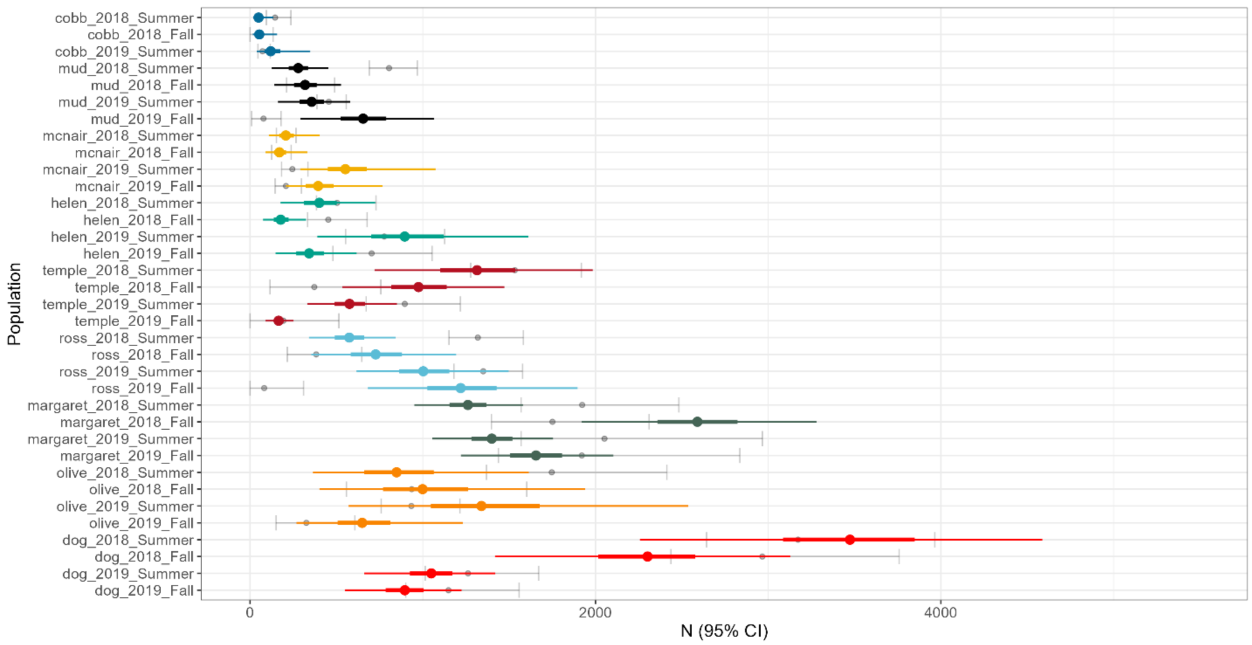


**Figure SM4**. Total abundance estimates from eDNA for all the populations and sampling periods. The colored thicker line represents the eDNA 50 % credible interval, and the colored thinner lines represent the eDNA 95 % credible intervals. The grey lines and dot represent the 95% credible interval and mean from the mark re-capture abundance estimates. eDNA estimates are produced from the fitted mass balance model (equation 8.1).

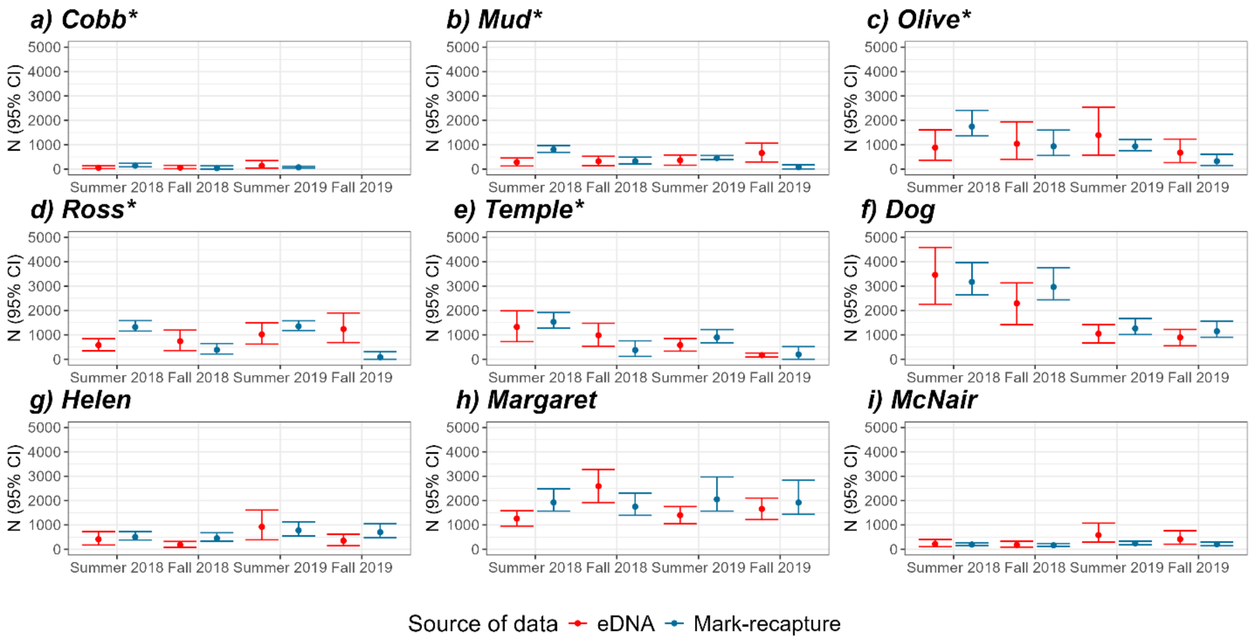


**Figure SM5.** Total abundance estimates from the eDNA mass balance model without sampling period specific *I0* (red) and mark-recapture (blue). eDNA estimates are produced from the fitted mass balance model (equation 8 in the main manuscript). The * mark lakes that were harvested. eDNA estimates are produced from the fitted mass balance model without sampling period specific eDNA generation coefficient (I0) (equation 8.1).

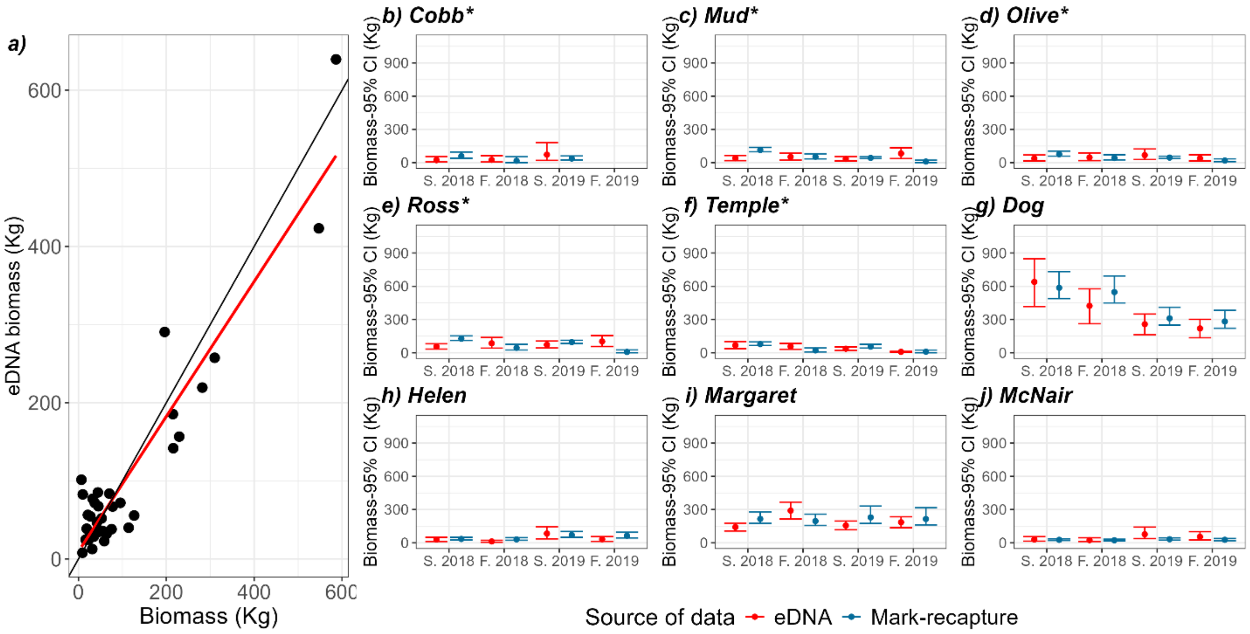


**Figure SM6.** Biomass estimated from the eDNA mass balance model and mark-recapture. In a) the x axis is the biomass estimated from the mark-recapture abundance estimates and the y axis is the biomass estimated from the eDNA mass balance model. The red line is the linear regression between the two and the black line is the 1 for 1 slope. In b) to j) the biomass estimated from the eDNA mass balance model (red) and the biomass estimated from the mark-recapture (blue). eDNA estimates are produced from the fitted mass balance model without sampling period specific eDNA generation coefficient (I0) (equation 8.1). The * mark lakes that were harvested.

### Appendix 6: Population estimates when not including a K estimate for each lake nor sampling period specific I0.

To know the importance of considering differences in eDNA disappearance rates (*k*) between lakes, a model without lake specific k was fitted. This model had lower performance with a bayes $R^2$ of 0.5 in comparison to a bayes $R^2$ of 0.69 for the mass balance model with lake specific k (Table 2). Consequently, the population estimates produced are less accurate (fig SM3).

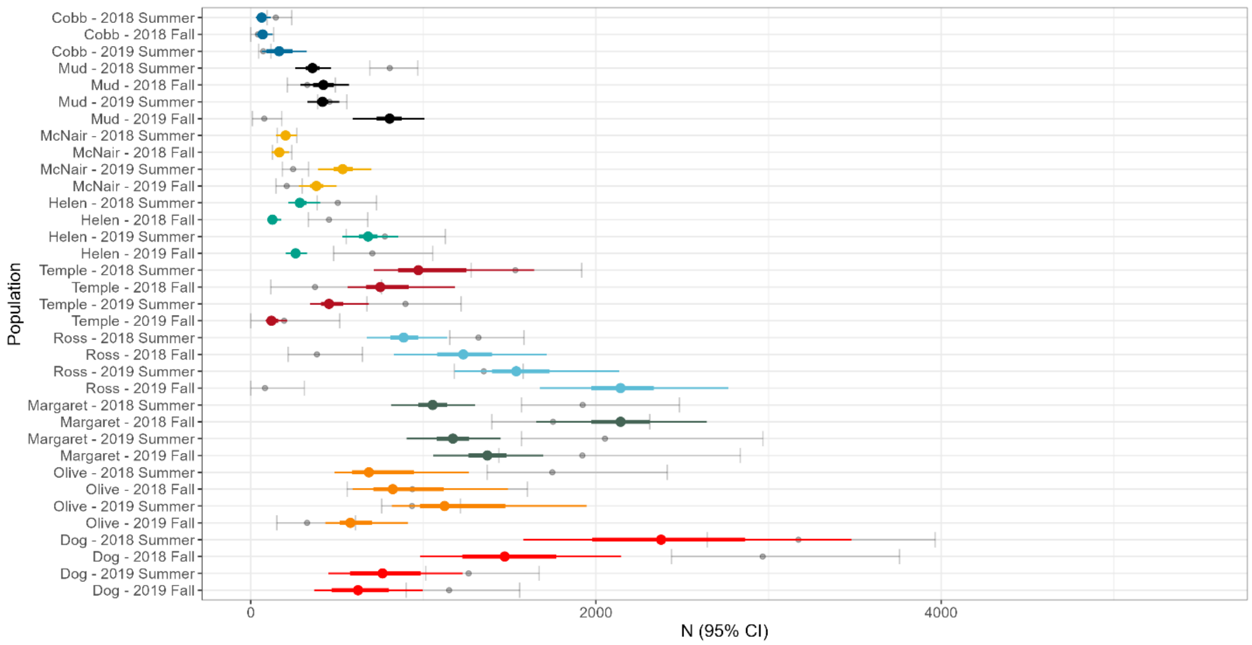


**Figure SM7**. Total abundance estimates from eDNA for all the populations and sampling periods. The colored thicker line represents the eDNA 50 % credible interval, and the colored thinner lines represent the eDNA 95 % credible intervals. The grey lines and dot represent the 95% credible interval and mean from the mark re-capture abundance estimates. eDNA estimates are produced from the fitted mass balance model (equation 8.1).

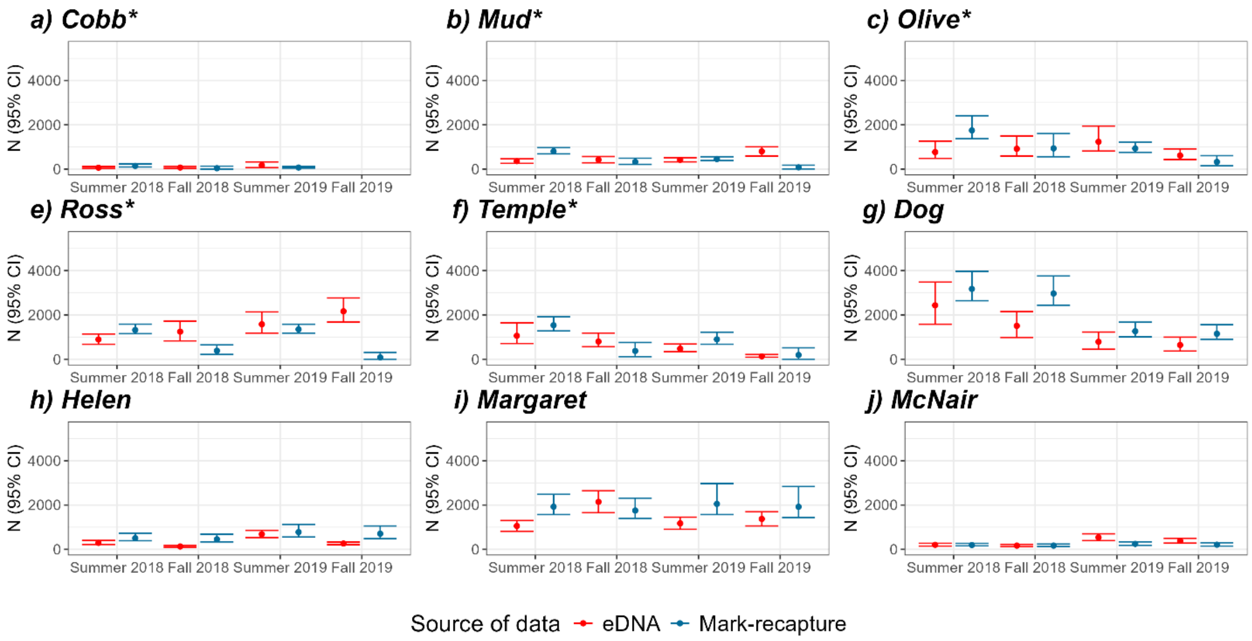


**Figure SM8**. Total abundance estimates from eDNA (red) and mark re-capture (blue). eDNA estimates are produced from the fitted mass balance model (equation 8), but with without a lake specific eDNA disappearance rate (*k*). The * mark lakes that were harvested. The error bars represent the 95% credible interval.

### Appendix 7: Bias testing with the mass balance model

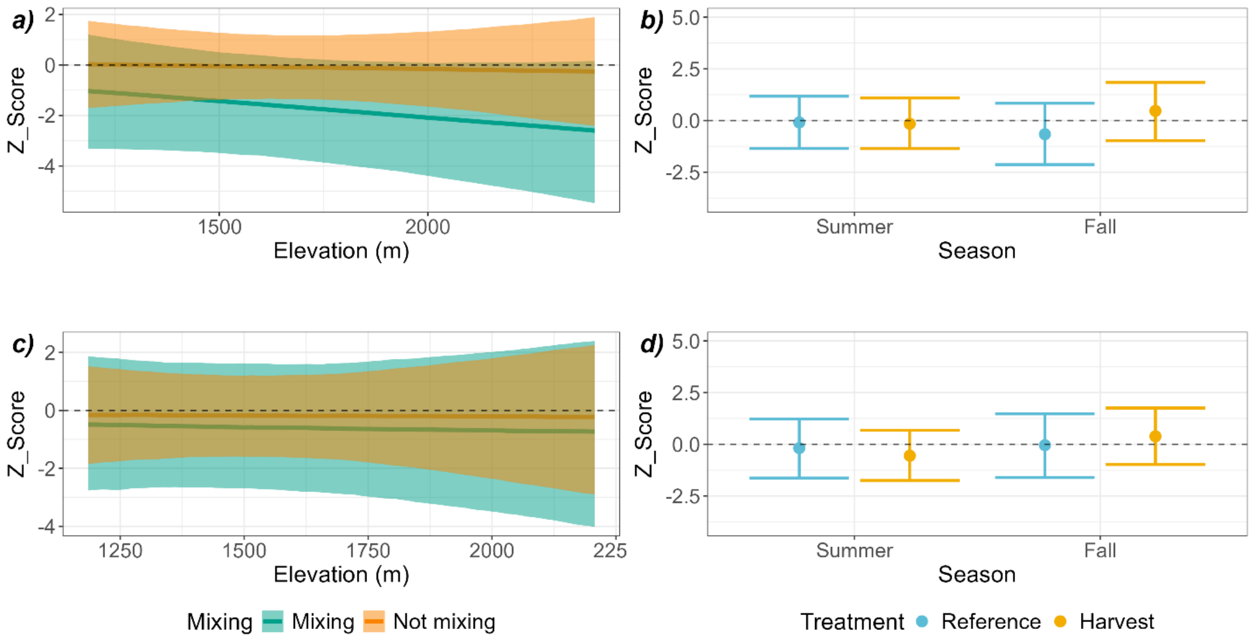


**Figure SM9.** Effect of thermal mixing, elevation and experimental treatment (harvested versus reference lake) on the eDNA mass balance model performance. The figure shows the marginal posterior predictions with 95 % credible interval. In a) and b) the model was fitted with all lakes while in c) and d) Helen Lake was removed to see if the trends observed were driven only by Helen Lake. The model parameter estimates and $R^2$ of each model is reported in Table 3. The z-score was calculated as the estimate from eDNA minus the estimate from mark-recapture divided by the standard deviation of the draws of the estimate from eDNA.

### Appendix 8: eDNA turnover time

To estimate the eDNA turnover rate, or the time for the eDNA to disappear from a lake if there was no more eDNA production, we took the summer (pre-harvest) eDNA concentration as the initial value and ran simulations for each and each year to estimate the number of days necessary for eDNA to degrade considering both:

$$[eDNA]_t = \alpha\, e^{-kt} - \frac{Q\,[eDNA]_{t-1}}{V}$$

Where $[eDNA]_t$ is the eDNA concentration at time $t$, $\alpha$ is the initial (summer) eDNA concentration, $Q$ is the outflow rate, $V$ is the epilimnion volume, and $k$ is the eDNA disappearance constant parameterised by the full mass balance model. Estimations were bootstrapped to include the uncertainty on $k$ and the eDNA concentration at $t-1$.

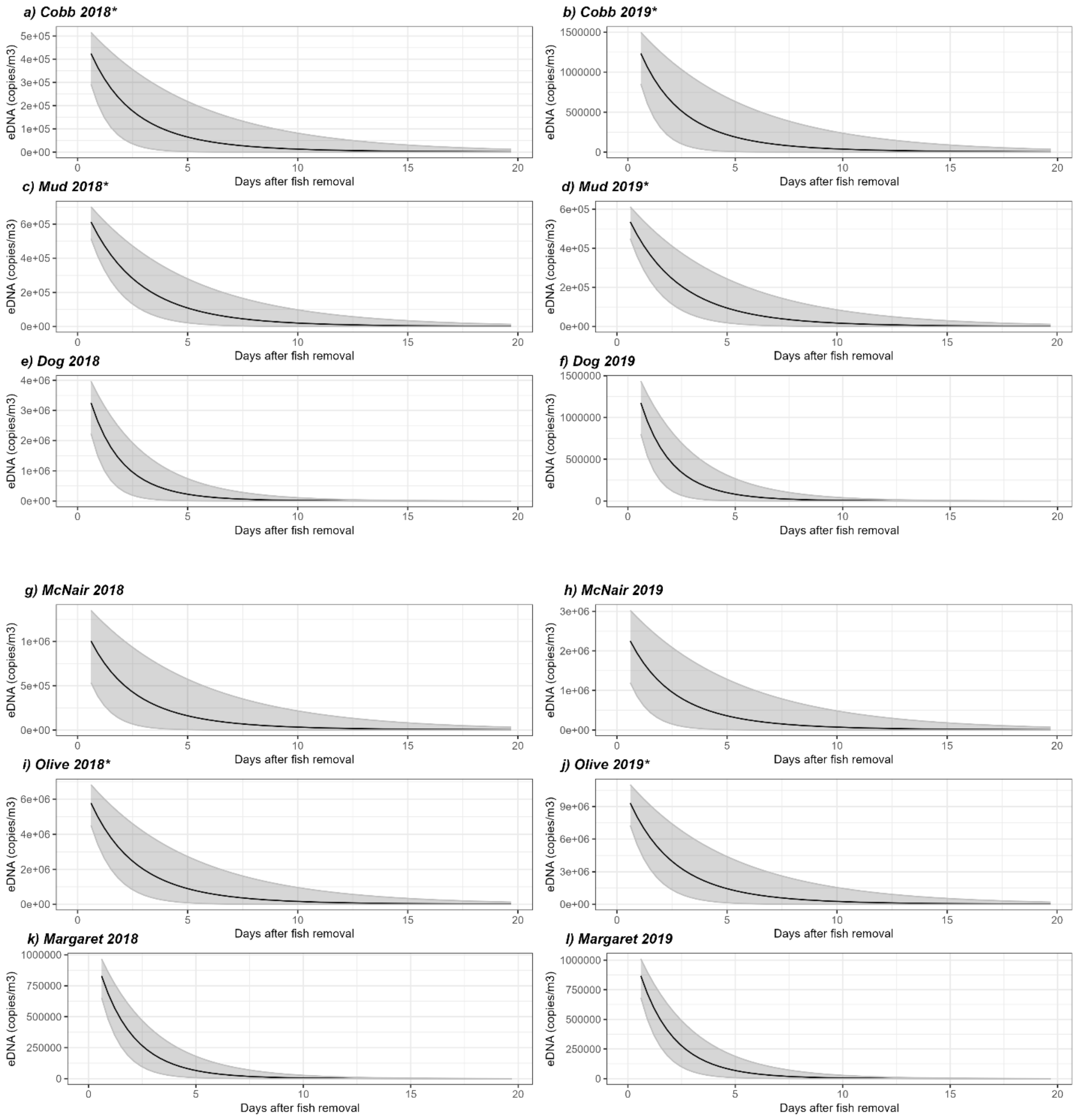

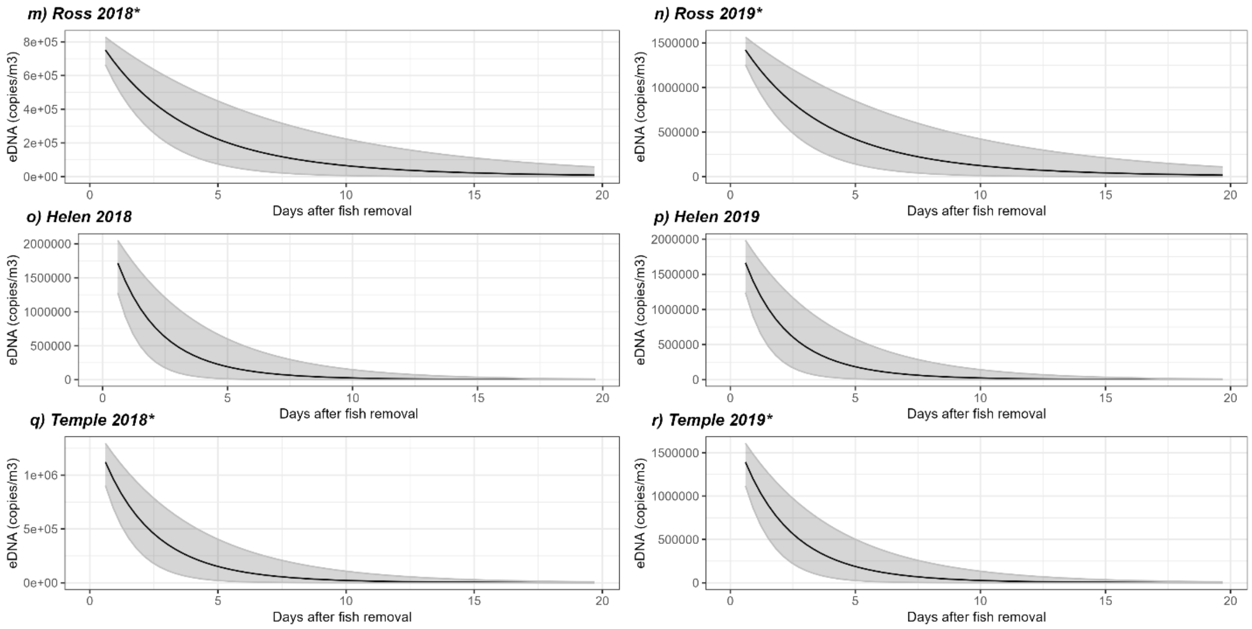


**Figure SM10:** Time for eDNA turnover. The grey area represents the 95% confidence interval, and asterix (*) stands for lakes that were harvested.
